# Comparative characterization of OncoPro™ and Wnt-Based media reveals distinct phenotypic and pharmacologic states in patient-derived tumor organoids

**DOI:** 10.64898/2025.12.13.693944

**Authors:** Sofie Seghers, Maxim Le Compte, Felicia Rodrigues Fortes, Jana Baroen, Jasper Ott, Geert Roeyen, Vera Hartman, Samir Kumar-Singh, Lenny Coppens, Michiel de Maat, Niels Komen, Sylvie Van Den Broeck, Jody Valk, Wiebren Tjalma, Jeroen M.H. Hendriks, Paul Van Schil, Gilles Van Haesendonck, Sofia L. J. Peeters, Hans Prenen, Christophe Deben

## Abstract

**Background:** Patient-derived tumor organoids (PDTOs) are strongly influenced by culture medium. We compared OncoPro™ (OP) Tumoroid Culture medium with conventional Wnt/R-spondin/noggin (Wnt) medium. This recently developed OP medium offers a standardized, serum-free alternative to Wnt-based formulations.

**Methods:** We compared OP and Wnt media across 36 PDTO lines from various malignancies (colorectal, pancreatic, breast, lung, gastric, gastroesophageal junction, biliary head- and neck and uknown primary), assessing establishment success. Selected PDTO models were subjected to downstream characterization, including morphological assessment and bulk transcriptomic profiling with comparison to public single-cell RNA sequencing reference datasets (n=11), whole-exome sequencing (WES) (n=9), and pharmacological response profiling to a 33-drug panel (n=3).

**Results:** Adaptation from Wnt medium to OP succeeded in 83.3% (15/18), whereas de novo establishment favored Wnt (33.3% vs 11.1%). Key oncogenic driver alterations were retained across matched organoid cultures, supporting preservation of tumor-relevant genomic features. Transcriptomic profiling confirmed preserved tumor-identity across media, while revealing different epithelial state programs: Wnt upregulated proliferation/stemness-associated genes (e.g. LGR5) and OP enriched adhesion-associated genes and inflammatory/TGF-β programs. In scRNA databases OP signatures preferentially mapped to malignant epithelial compartments in pancreatic cancer, whereas Wnt signatures were linked to non-malignant epithelium. Similarly, in colon cancer OP signature mapped predominantly to the malignant epithelial compartments. Drug (n=33) screening in pancreatic- and colorectal cancer PDTOs (n=3) demonstrated consistent medium-dependent shifts: Wnt-grown PDTOs were globally more sensitive in the screened subset, particularly to MAPK-axis inhibitors and apoptosis-sensitizers, while OP-grown PDTOs exhibited relative resistance.

**Conclusions:** Culture medium composition is a key determinant of PDTO phenotype, transcriptome and drug sensitivity. Wnt medium was associated with drug-sensitive states, whereas OP medium was associated with adhesion- and inflammatory-related programs, relative resistance in the screened subset of 33 drugs and closer alignment with malignant epithelial programs in the analyzed pancreatic / colorectal cancer single-cell atlases.

## Introduction

Patient-derived tumor organoids (PDTOs) are three-dimensional (3D) cell cultures originating from patient tumor samples, in which cancer cells self-assemble into clusters that maintain the original tumor’s histological and molecular characteristics. These PDTOs exhibit self-renewal properties and closely mimic the epithelial heterogeneity of the tumor and its complexity ex vivo, providing a physiologically relevant model for studying cancer biology, drug response and personalized medicine [1–5]. Central to the success of PDTO cultures is the composition of the culturing medium, which plays a crucial role in maintaining the viability and functionality of these organoids. The optimization of PDTO culture medium has been an important point of research efforts, with emphasis on the inclusion of seemingly essential growth factors such as Wnt, Noggin and R-spondin [6, 7]. These factors are pivotal for sustaining the stemness and self-renewal capacity of tumor cells within the organoid structure [8]. Specifically, Wnt signaling drives stem cell proliferation and maintenance, while R-spondin enhances Wnt activity by binding to the LGR5 receptor, a marker of epithelial stem cells. Noggin, on the other hand, inhibits bone morphogenetic protein signaling, maintaining tumor cells in an undifferentiated state [9]. Together, these components create a microenvironment that mimics the niche required for tumor cell survival and growth. Recently, Thermo Fisher Scientific introduced the serum-free Gibco™ OncoPro™ Tumoroid Culture medium (OP), a novel alternative to existing formulations without the presence of Wnt-3A, R-Spondin, noggin or small molecule inhibitors. This medium provides broad freedom to operate, facilitating use in both academic research and commercial development, while ensuring broad accessibility to essential growth factors, offering standardized formulations for multiple cancer types, including lung, colorectal, pancreatic, head and neck, endometrial and triple-negative breast cancer [10].

In this study, we performed a comparison of the OP medium with conventional Wnt/R-spondin/noggin–based formulations (hereafter referred to as Wnt medium) across multiple tumor types. We assessed de novo PDTO establishment rate, adaptation from existing Wnt cultures, and effects on growth dynamics, morphology, lineage marker preservation and oncogenic drivers. Bulk RNA-sequencing (RNAseq) revealed that OP and Wnt media impose distinct transcriptional states. By integrating bulk transcriptomes with public single-cell RNA-sequencing (scRNAseq) datasets from pancreatic ductal adenocarcinoma (PDAC) and colorectal cancer (CRC), we assessed how these medium-induced states correspond to patient epithelial tumor programs. We further assessed functional consequences of medium choice on drug response using high-content phenotypic screening, identifying key differences in response depending on the therapeutic class. This study provides critical insights into the advantages and limitations of OP and highlights the importance of medium composition for assessing ex vivo drug response on PDTOs.

## Materials and Methods

All used reagents and resources (including cell culture reagents, antibodies and drugs) can be found in **Table S1: Resources and reagents.**

### Patient samples

Tissue resection fragments were procured from patients undergoing surgery or thoracocentesis/paracentesis at the Antwerp University Hospital (UZA), with written informed consent obtained from all participants. This study was conducted as a pan-cancer organoid platform; therefore, tumor specimens from multiple anatomical sites were eligible for inclusion and the final tumor-type distribution consists out of pancreatic, colorectal, gastric, gastro-esophageal junction, lung, biliary tract, head-and neck, breast cancer and a carcinoma of unknown primary. The study protocols were approved by the UZA Ethical Committee (ref. 14/47/ 480 for PDAC; ref. 2022-3470 for gastro-intestinal tumors; ref. 20/08/090 for head and neck tumors; ref 2023-5364 for other cases). Human biological specimens were obtained from Biobank@UZA (Antwerp, Belgium; ID: BE71030031000), supported by the Belgian Virtual Tumorbank under the National Cancer Plan.

### Patient-derived tumor organoid establishment and culture

Tumor resection specimens or biopsies were preserved in collection medium (**Table S2: media**) at 4°C and transported on ice for processing within 24 hours for PDTO culture. Upon arrival, the tumor fragments were dissected on ice to eliminate residual connective tissue and cut into approximately 1-2 mm^2^ pieces. Subsequently, the tissue fragments underwent a wash with ice-cold phosphate-buffered saline (PBS) followed by enzymatic digestion at 37°C for 1 hour using digestion medium (**Table S2: media**). After digestion, the cell suspension was filtered through a 100 μm filter and the filtrates were centrifuged at 1500 rpm for 5 minutes. The resulting cell pellet was resuspended in ice-cold Cultrex growth factor reduced basement membrane extract type 2. Small droplets of 20 μL were plated and inverted for 30 minutes at 37°C to allow solidification. These droplets were then overlaid with tumor-specific medium. To prevent microbial contamination, 1x Primocin was added to the culture medium for the initial two weeks. The specific composition of the medium can be found in **Table S2: media**.

For passaging, PDTOs were dissociated into single cells using TrypLE Express. Cryopreservation was performed by harvesting 3-day-old PDTOs with Cultrex Harvesting Solution and freezing them in Recovery Cell Culture Freezing Medium. Successful PDTO establishment was defined as sustained expansion beyond passage 5 with exponential growth sufficient for cryopreservation and preservation of genomic features consistent with malignancy of the original tumor tissue. Additionally, samples were screened for Mycoplasma contamination using the MycoAlert Mycoplasma Detection Kit.

Model fidelity was assessed using patient-matched germline fingerprinting, retention of tumor-type-relevant oncogenic driver alterations by WES, lineage marker expression, and exclusion of cultures with evidence of non-malignant epithelial overgrowth.

### Transfer to OncoPro™ Tumoroid Culturing medium

The OP medium was prepared according to the manufacturer’s specifications. The specific composition of the medium utilized for each tumor type can be found in **Table S2: media.** Initially, the transfer of the PDTOs to OP was carried out using an ‘immediate’ transition method, involving direct transfer after passaging into the new medium. Subsequently, a ‘gradual’ transition approach was employed, wherein 25% of the OP was incrementally introduced after each passage (i.e., 25%, 50%, 75%, 100). A visual representation can be found in **Fig. S1**. Successful adaptation to OP medium was defined as sustained growth for at least two consecutive passages in OP with clear exponential expansion.

### Drug screening

Drug screening on 3D organoids was performed at the DrugVision.AI automated screening facility of the University of Antwerp, Belgium, using a prevalidated drug screening pipeline for which a detailed protocol is available in the Journal of Visualized Experiments [11]. Briefly, established organoid lines were expanded in extracellular matrix (ECM) domes (Cultrex type 2). Next, 3-day-old organoids were harvested from ECM drops using the Cultrex Organoid Harvesting Solution, collected in a 15 mL tube coated with 0.1% bovine serum albumin (BSA)/PBS, washed, and resuspended in medium. Next, the number of organoids was quantified by adding 5 µL of the organoid solution to 45 µL of medium in a 384-microplate well. A whole-well brightfield image was captured using the Tecan Spark Cyto and the number of organoids was counted label-free using Orbits^®^ (Orbits Oncology). Next, the organoid solution was diluted in full medium (without NAC or Y-27632) supplemented with 4% Cultrex at a concentration of 4000 organoids / mL. Next, 50 µL (200 organoids) of this solution was dispensed into each well of a 384-well ultra-low attachment microplate (Corning, #4588) using the OT-2 pipetting robot (Opentrons) in a cooled environment. Thereafter, the plate was centrifuged (100 rcf, 30 s, 4 °C) to ensure that all organoids are in the same z-plane and incubated overnight at 37 °C.

Drugs and fluorescent reagents were added to the plate using the Tecan D300e Digital Dispenser. Compounds were screened using a 4-point logarithmic dose titration within predefined concentration ranges routinely applied in our platform.

– High range (100- 20000 nM, log scale (100 nM, 271 nM, 737 nM, 20000 nM)): 5-Fluorouracil (5-FU), cisplatin, oxaliplatin.
– Middle range (100–3000 nM, log scale (100 nM, 311 nM, 966 nM, 3000 nM)): A-770041, afatinib, anlotinib, ASTX029, axitinib, barasertib, birinapant, buparlisib, cetuximab, erlotinib, everolimus, gefitinib, ibrutinib, MK2206, MRTX-1133, navitoclax, neratinib, olaparib, palbociclib, rucaparib, selumetinib, simvastatin, trametinib, venetoclax, WH-4-023, ZM-44739.
– Low range (1–100 nM, log scale (1 nM, 4.64 nM, 21.5nM, 100 nM): docetaxel, gemcitabine, paclitaxel, SN-38.

Staurosporine was used as positive control. No medium was changed during the experiment. Compounds were dissolved in DMSO, except for cetuximab, oxaliplatin and cisplatin, which were dissolved in 0.66% Tween-20 in PBS for dispensing with the D300e Dispenser. Brightfield and green fluorescence whole-well images (4x objective) were captured at T0, T60 and T120 using the Tecan Spark Cyto set at 37 °C / 5% CO2 for 5 days. This study included two technical and two biological replicates per patient, with biological replicates defined as PDTOs being cultured in separate wells for ≥2 passages. Drug screening experiments were performed only after cultures had stabilized in their respective media (≥2 passages after reaching 100% OP or Wnt conditions). Passage numbers can be found in **Table S3: passages**.

### Image and data analysis

Images were analyzed with the Orbits^®^ label-free organoid detection module [12]. The Normalized Organoid Growth Rate (NOGR) drug response metric was used for all downstream data analysis, which is described in detail in Deben et al., 2024 [13]. Based on the NOGR, the drug effects can be classified as: >1, proliferative effect; = 1, normal growth as in negative control; = 0, complete growth inhibition; = −1, complete killing as in positive control. In addition, we also calculated the Area Over the Curve (AOC) and normalized it to the maximal area. This ratio provided a value between 0 and 1 to reflect no effect and complete effect of the drug, respectively.

### RNA sequencing

For RNAseq, OP- and Wnt-cultured PDTOs were grown side by side and processed in parallel at matched time points to ensure direct comparability. For each condition, PDTOs were cultured in three independent wells, generating three replicates per medium. Sequencing was performed only after cultures had stabilized in their respective media (≥2 passages after reaching 100% OP or Wnt conditions) (**Fig. S1**). Passage numbers can be found in **Table S3: passages**. The cohort included CRC (n=4), PDAC (n=3), gastroesophageal junction carcinoma (GEJC) (n=3), carcinoma of unknown primary (CUP) (n=1), and head-and neck squamous cell carcinoma (HNSCC) (n=1). Subsequently, RNA extraction was performed using the RNeasy blood midi kit according to the manufacturer’s instructions. GEJC025WNT_rep3 and CRC018WNT_rep3 were removed due to poor sample quality. The concentration of the extracted RNA was assessed using the NanoDrop ND-1000. Following assessment, samples were frozen at −80 °C and transferred to the Genomics Core Leuven for transcriptome sequencing using Lexogen QuantSeq 3’ FWD library preparation kit for Illumina on a Hiseq400 SR50 line with a minimum of 2M reads per sample. Downstream analysis and plotting (clustered heatmaps, uniform manifold approximation and projection (UMAP), Volcano plot, gene set enrichementenrichment analysis (GSEA)) were performed using the Omics Playground tool (Big Omics Analytics [14]) and python.

### Whole exome sequencing

For WES, OP- and Wnt-cultured PDTOs from nine patients were expanded side by side and harvested in parallel at matched time points, enabling direct comparison between culture conditions. The cohort included CRC (n=4), PDAC (n=3), GEJC (n=1), and HNSCC (n=1). Sequencing was performed only after culture stabilization, as defined for the RNA-sequencing experiments; corresponding passage numbers are provided in **Table S3**: **passages**. One replicate, consisting of a dry pellet, was included per PDTO line. Parental tumor material consisted of formalin-fixed paraffin embedded (FFPE) rolls. PDTO dry pellets and FFPE rolls were submitted to GeneWiz by Azenta for library preparation and exome capture using the Twist Exome 2.0 kit. Parental tumor samples were sequenced to 10 Gb, corresponding to approximately 30M read pairs, whereas PDTO samples were sequenced to 5 Gb, corresponding to approximately 15M read pairs. Resulting in a final WES dataset of 24 samples, comprising parental tumor tissue (n=6), OP-cultured PDTOs (n=9), and Wnt-cultured PDTOs (n=9). The CRC013 parental FFPE sample was excluded from mutation analysis because insufficient DNA yield resulted in poor sequencing quality metrics reported by Genewiz.

#### Variant calling

Raw reads were processed using nf-core/sarek (v3) [15, 16] with the GRCh38/hg38 reference genome. Briefly, reads were quality-assessed with FastQC [17] and aligned using BWA-MEM2 [18]. Somatic variants were called with GATK Mutect2 [19, 20] in tumor-only mode, with panel-of-normals filtering. Variant filtering applied gnomAD [21] and 1000 Genomes [22] allele frequency thresholds, GATK LearnReadOrientationModel, and strand-bias filters (OPTIMAL_F_SCORE, FDR=0.05). Minimum PASS thresholds: MQ≥20, base quality≥18, tumor LOD≥3.0, normal LOD≥2.2.

#### Clonal mutation recovery

Organoid-derived variants suppressed as germline or normal artifact artefacts by Mutect2 were systematically recovered. Parental PASS variants were cross-referenced against organoid VCFs: suppressed organoid calls overlapping a parental PASS site were reclassified as clonal. VCF parsing used vcfR [23]; genomic overlaps were resolved with GenomicRanges [24]. Functional annotation was applied via VariantAnnotation [25] and Bioconductor infrastructure [26]. Mutations were classified as Clonal (shared parental/organoid), Organoid_Acquired (organoid-only), or Parental_Specific (parental-only).

#### Mutational signatures

Single-base-substitution (SBS) trinucleotide profiles were constructed from somatic PASS variants. COSMIC v3.3 SBS signatures were fitted using MutationalPatterns [27] with non-negative least-squares decomposition; signatures contributing <5% were excluded.

#### Statistical analysis and visualization

All analyses were performed in R or Python [28]. Mutation co-occurrence matrices were generated with ComplexHeatmap [29]; additional graphics with ggplot [30]. For mutation plotting, WES variants were filtered to genes listed in the COSMIC Cancer Gene Census Tier 1 (CGC, v99). Only somatic single-nucleotide variants classified as Missense_Mutation, Nonsense_Mutation, or Splice_Site with a variant allele frequency (VAF) ≥ 0.05 were retained. For each patient, three sample types were analysed: parental tumour (T, if available), OP-derived organoids (O), and WNT-treated organoids (W). Mutations were aggregated per gene–sample pair; if multiple variants were detected in the same gene and sample, the entry was labelled Multi_Hit and the maximum VAF was used. An OncoPrint-style mutation landscape was generated in Python using matplotlib. Each cell represents one gene–sample combination; cell height is proportional to the VAF of the dominant variant, providing a visual estimate of clonal abundance. Positions with zero or missing sequencing depth in a sample were flagged with an asterisk (*) if the same gene carried a mutation in at least one other sample from the same patient, indicating ambiguous absence of the variant rather than confirmed wild-type status.

### Integrated PDAC and CRC single cell atlases

For PDAC, the epithelial sub-atlas from the PDAC atlas was downloaded from the UCSC Cell Browser at https://cells.ucsc.edu/?ds=pdac-atlas and used without further modifications [31]. The CRC atlas was downloaded from https://doi.org/10.6084/m9.figshare.25323397 [32]. Using the provided annotations ‘Epi’ and ‘Malignant Cells’ under ParentalCluster, we generated an epithelial sub-atlas containing epithelial cells derived from CRC tumor, healthy, and polyp patient samples. Highly variable genes were identified using Scanpy’s highly_variable_genes function (min_mean=0.0125, max_mean=3, min_disp=0.5). Data were scaled (max_value=10) and principal component analysis was performed. Patient batch effects were corrected using Harmony on the PCA embeddings (theta=2.0, max_iter=20). Neighborhood graphs were computed using Harmony-corrected embeddings (n_neighbors=10, n_pcs=50), followed by UMAP dimensionality reduction. Leiden clustering was performed at resolution 0.2 to identify cell subpopulations. Differential gene expression analysis was conducted using the Wilcoxon rank-sum test to identify cluster-specific marker genes, excluding mitochondrial genes from visualization. Custom gene sets were established based on the top 50 upregulated genes in OP medium or Wnt-based medium, independently for PDAC and CRC samples (analyzed via differential expression BigOmics).

Gene set activity scores were computed at single-cell resolution using the AUCell algorithm [33], as implemented in the decoupler v2 Python package [34]. AUCell assigns each cell an Area Under the Curve (AUC) score by ranking all expressed genes by their log-normalised expression level and calculating the fraction of each gene set’s members that falls within the top-ranking genes (default: top 5% of the transcriptome). Scoring was applied to log-normalized expression values (UMI counts normalized to 10,000 reads per cell followed by log₁p transformation). Gene sets with fewer than five detected target genes per cell were excluded (tmin = 5). To limit peak memory usage, scoring was performed sample-by-sample.

### Cytokine assay

To quantify cytokines, present in the medium, we collected supernatants from CRC018 grown in both Wnt and OP medium. At T0 we collected samples of freshly prepared Wnt and OP CRC medium (OP with (+BSA) and without (-BSA) BSA) to evaluate baseline cytokines. CRC018 organoids were then seeded at ∼200 organoids per well in a 384-well ultra-low-attachment plate (Corning #4588) in their respective media. Supernatants were sampled at 24 hours (T24) and 120 hours (T120) post-seeding and frozen at –80 °C until analysis. Of note, all cytokine experiments were analyzed in triplicate, apart from the TGF-β total and active, of which there was only one replicate.

Levels of 49 cytokines were measured in serum samples using U-plex (K15067M & K151XWK) and V-plex (K15198D) panels from Meso Scale Discovery (MSD, MD, USA), according to the manufacturer instructions. Included cytokines can be found in supplementary **Table S4: cytokines.** Measurements were performed in randomized batches and read on the QuickPlex SQ 120 (MSD). Briefly, 96-well plates of the U-plex panels were coated with a capturing antibody linked to a linker for one hour. The vascular injury panel (V-plex) was washed before use. All plates were then washed three times with MSD wash buffer. Samples were incubated for two hours (except for the V-plex panels and BDNF panel, which had one hour of incubation), following incubation the plates were washed three times. Detection antibody with a sulfo-tag was added and after another one-hour incubation step (two hours for the angiogenesis panel), plates were washed and read with MSD reading buffer on the QuickPlex SQ 120. Throughout the statistical analyses, cytokine values below the detection range were recorded as 1x the lower limit of detection (LLOD) and values above the detection range were recorded as 1x the upper limit of detection.

### Immunohistochemistry

Immunohistochemical staining was performed on a Dako Omnis platform (Agilent Technologies) using EnVision Flex detection reagents and DAB visualization, followed by hematoxylin counterstaining. TTF-1 staining was performed after 30 min heat-induced epitope retrieval in high-pH buffer, followed by 20 min primary antibody incubation (clone SPT24, Leica, 1:400) and a 10 min mouse linker step. p63 staining was performed after 30 min retrieval in low-pH buffer with 25 min primary antibody incubation (clone DAK-p63, Agilent/Dako, ready-to-use) without linker. CK5/6 staining was performed after 30 min retrieval in high-pH buffer with 13 min primary antibody incubation (clone D5/16 B4, Agilent/Dako, ready-to-use) followed by a 10 min mouse linker step. Napsin A staining was performed after 30 min retrieval in high-pH buffer with 20 min primary antibody incubation (clone MRQ60, Cell Marque, 1:500) without linker. Appropriate external control tissues were included in each run and staining results were evaluated by a certified pathologist.

### Statistical analysis

Confidence intervals for establishment and adaptation success rates were calculated using Wilson binomial 95% confidence intervals, which are appropriate for proportions derived from small sample sizes.

Statistical analysis of bulkRNAseq was conducted using BigOmics, a bioinformatics platform designed for the integrative analysis of multi-omics data [14]. Raw gene expression data were normalized to correct for batch effects and systematic biases. Depending on the dataset, normalization methods such as quantile normalization or Transcripts Per Million (TPM) were applied to ensure comparability across samples. Differential expression analysis was performed using DESeq2, edgeR and limma to identify genes significantly regulated by the culture medium. GSEA was used to identify enriched biological pathways, leveraging multiple integrated databases within BigOmics, including Hallmark dataset. To further investigate differential drug sensitivity based on transcriptomic profiles, the BigOmics drug connectivity module with the qCTRPv2 and GDSC databases was used.

Data visualization was carried out using various tools. BigOmics for bulk RNAseq data (see above). BioRender was employed to illustrate compound overviews. Other images were created in Python.

## Results

### Patient cohort and sample overview

A total of 36 patients were included in this study (**Table S5: clinical characteristics**). The majority had digestive malignancies, including PDAC (*n* = 12), CRC (*n* = 8), CCA( *n* = 4), GEJC (*n* = 2), and gastric cancer (*n* = 1). Other tumor types comprised HNSCC (*n* = 3), breast cancer (*n* = 2), non-small cell lung cancer (NSCLC, *n = 2)*, renal cell carcinoma (*n* = 1) and carcinoma of unknown primary (CUP, *n* = 1). Most tumors were adenocarcinomas (*n* = 32) and most patients were treatment-naïve (*n* = 25). Specimens were mainly obtained from surgical resections or biopsies, with four derived from malignant effusions.

### De novo establishment and adaptation success rates

We evaluated the de novo establishment from fresh tumor material (n = 18) or adaptation capacity (n = 18), defined as the successful transfer from Wnt-based medium to OP culture medium, of 36 PDTO lines of various tumor types. The overall culturing success rate in OP medium, stratified by tumor subtype, pretreatment status, cryopreservation status and micro-satellite instable (MSI) status, is shown in **Figure 1A**. Particularly, squamous cell carcinoma (SCC) samples and MSI-positive tumors displayed the highest success rates. These observations should, however, be interpreted with caution given the limited number of samples in these subgroups. NSCLC-derived samples were not included here due to recurrent non-tumor epithelial overgrowth as described below. Among the transitioned lines, 83% (15/18; 95% CI 60.8–94.2%) successfully adapted to OP (**Table 1**, **Fig. 1B**). Three PDTO lines tolerated an immediate transition to 100% OP with minimal growth decline, whereas others initially engrafted but then underwent growth arrest and cell-yield reduction, prompting adoption of a gradual OP stepwise transition (25% - 50% - 75% - 100%) over multiple passages (see methods). This gradual method restored growth in most lines, though some PDTO lines (e.g. PDAC060, PDAC087 and GIASC001) still failed upon first passage in 100% OP.

**Figure 1:**
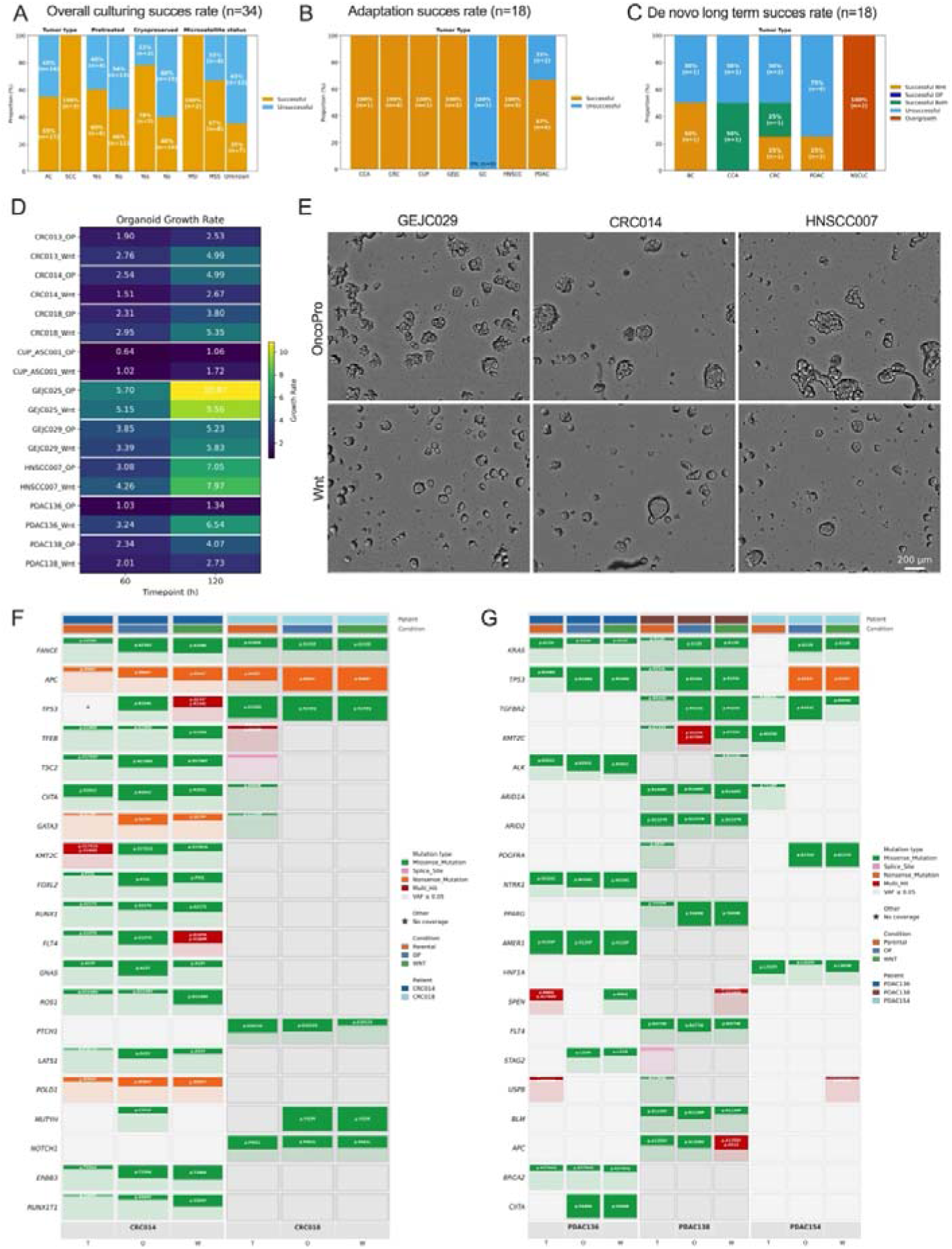
Culture medium influences organoid establishment and growth behavior across tumor types. **(A)** Overall organoid culturing success rate across tumor types (n = 34), stratified by pretreatment status, cryopreservation status, and microsatellite instability (MSI) status. Non-small cell lung cancer (NSCLC) were removed due to the overgrowth of non-malignant cells. Orange bars signify successful samples, blue bars unsuccessful samples; **(B)** Adaptation success rate in paired organoids transitioned from Wnt to OP medium (n = 18). **(C)** De novo long-term culture success rates by tumor type, indicating whether organoids were stably maintained in Wnt medium, OP medium, both, or neither (n = 18). None of the PDTOs were only successful in OP; Color of bars represent long term success status (orange = successful only in Wnt; dark blue = successful only in OP; light blue = unsuccessful; red = non-malignant overgrowth); **(D)** Organoid growth rates measured at 60 h and 120 h demonstrate mainly patient specific growth rates; **(E)** Representative brightfield images of paired PDTOs (GI029, CRC014, HNSCC007) cultured in OP versus Wnt medium illustrating distinct morphological phenotypes from matching passages; (**F-G**) Mutation landscape of top 20 COSMIC CGC genes across patient-derived samples in respectively CRC and PDAC. Each row represents a cancer gene (COSMIC CGC v99); each column represents an individual sample, grouped by patient (alternating grey bands) and annotated by sample type (Condition bar) and patient identity (Patient bar). Coloured rectangles indicate detected somatic mutations; cell height is proportional to the VAF of the dominant variant. A semi-transparent outline of the full cell height is shown for mutations with VAF < 0.05 to indicate low-frequency variants. An asterisk (*) marks samples with absent or insufficient sequencing coverage (total read depth = 0) at a locus where a mutation was detected in at least one other sample from the same patient. Mutation types are colour-coded as indicated in the legend. Only Missense_Mutation, Nonsense_Mutation, and Splice_Site variants are shown. <u>Abbreviations</u>: AC, adenocarcinoma; BC, breast cancer; CCA, cholangiocarcinoma; CUP, carcinoma of unknown primary; CRC, colorectal cancer; GC, gastric cancer; GEJC, gastro-esophageal junction carcinoma; GI, gastro-intestinal tumor; h, hour; HNSCC, head and neck squamous cell carcinoma; MSI, microsatellite instable; OP, OncoPro™ Tumoroid Culturing medium; PDAC, pancreatic ductal carcinoma; PDTO, patient-derived tumor organoid; SCC, squamous cell carcinoma; VAF, variant allele frequency Wnt, Wnt-supplemented medium

**Table 1:**
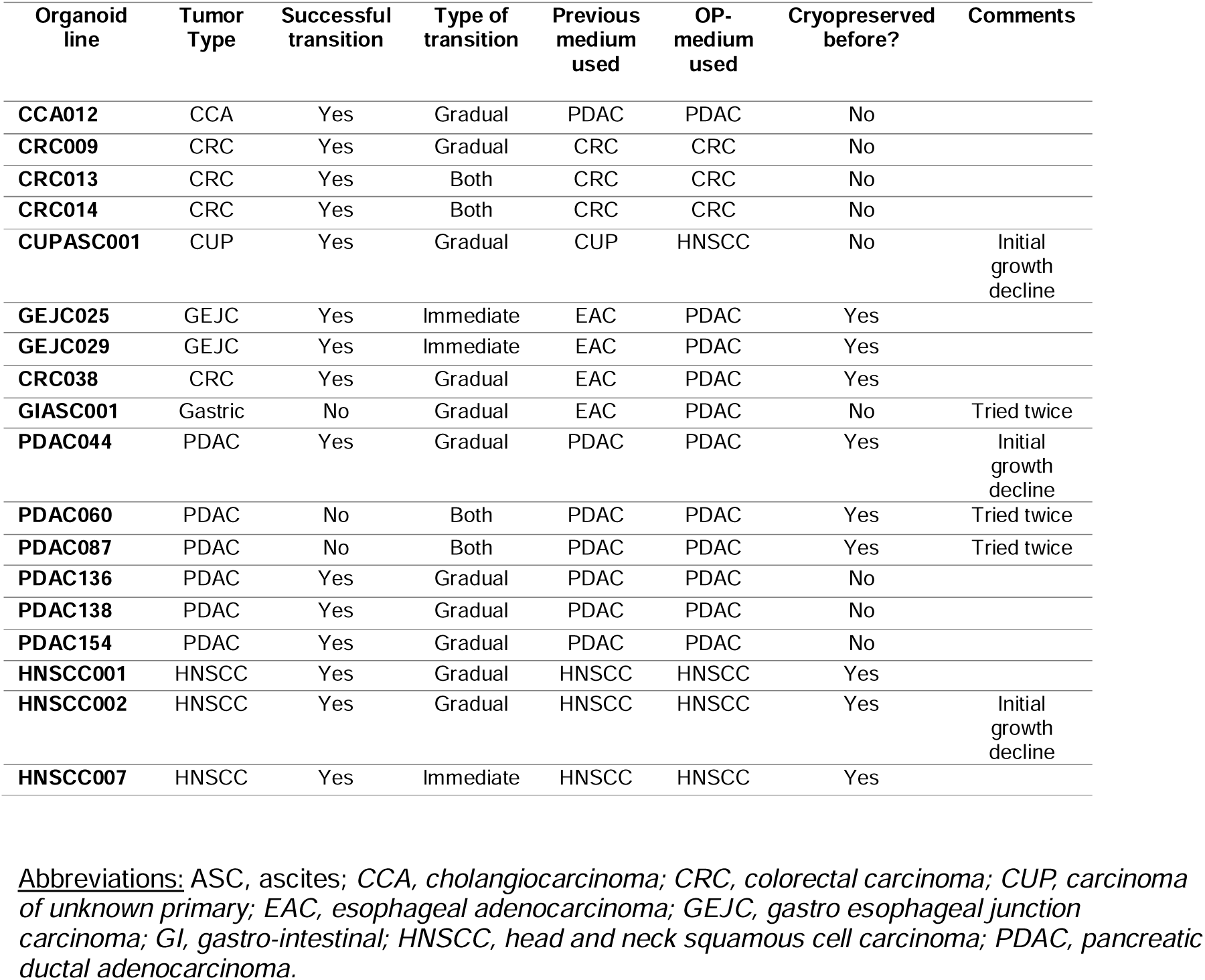
Overview of transition to OncoPro™ *tumoroid culture medium*.

De novo establishment was more efficient in Wnt medium (33.3%, 6/18 (95% CI 16.3–56.3%)) than in OP (11.1%, 2/18 (95% CI 3.1–32.8%)). Two lines survived in both media, but none exclusively in OP (**Table 2**, **Fig. 1C**). Interestingly, OP-established PDTOs frequently demonstrated superior early expansion, with seven lines showing larger diameters and cystic morphologies at the first passage in OP (vs. three favoring Wnt medium and six with no clear preference) (**Fig. S2A, table 2**). However, this early increase in organoid size did not translate into a higher long term establishment success rate.

**Table 2:**
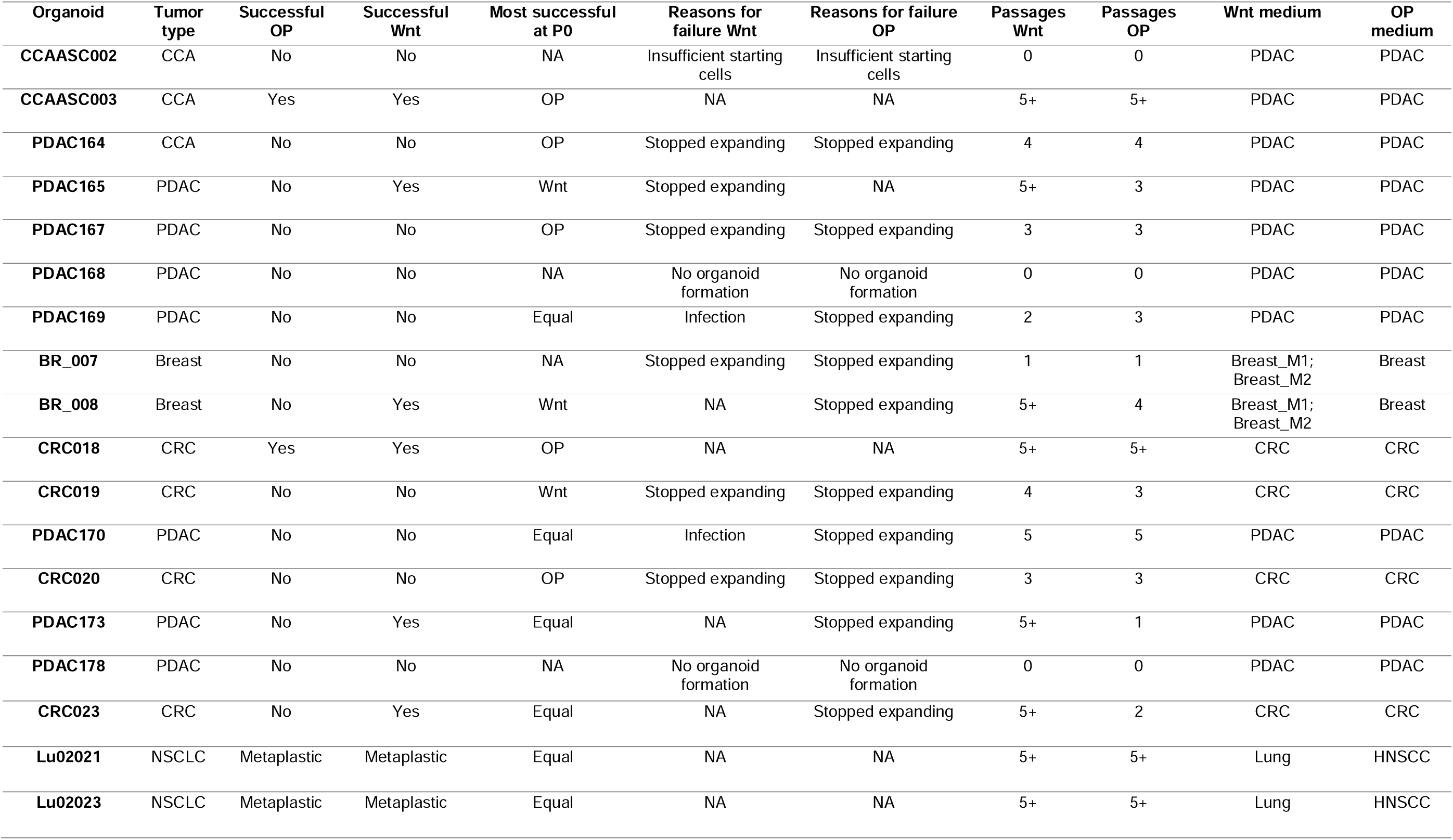
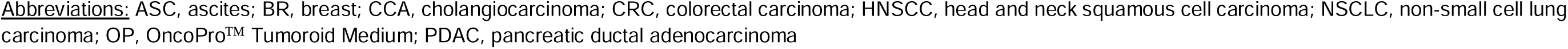
Overview of establishment of PDTOs starting from OncoPro™ *tumoroid culture medium*.

Additionally, given the well-described establishment challenges for pure NSCLC PDTOs [35], we evaluated whether OP medium could support tumor-specific outgrowth from NSCLC resections and endobronchial guided ultrasound fine needle biopsy (EBUS-FNA) samples. Although PDTOs emerged rapidly, immunohistochemical profiling [36] revealed that these cultures did not retain the marker signatures of their parental tumors. In Lu_02_021, the parental tumor exhibited a SCC phenotype (TTF-1−, p63+, CK5/6+, Napsin A−). In contrast, both the OP- and Wnt-derived PDTO cultures displayed a clear shift to TTF-1 positivity with heterogeneous p63 expression and retained CK5/6 positivity, while remaining Napsin A−, indicative of non-malignant overgrowth. Similarly, Lu_02_023, derived from a lung adenocarcinoma (expected TTF-1+/Napsin A+ and p63−/CK5/6−), produced organoid cultures that were TTF-1+ but Napsin A−, and displayed aberrant p63 and CK5/6 positivity, again indicative non-malignant overgrowth (**Fig. S2B)**. These NSCLC cultures were therefore excluded from downstream OP-versus-Wnt comparisons and are presented as a model-specific limitation, illustrating that OP medium did not overcome the known challenge of non-malignant epithelial overgrowth in NSCLC organoid establishment.

A subset of 11 organoid lines from various tumor types (CRC, PDAC, GEJC, HSNCC and CUP) was selected for further in-depth characterization and comparison between both culturing conditions. Growth rates were largely comparable between OP and Wnt conditions for most PDTOs, with no consistent growth advantage across media. Instead, differences in growth rates were generally patient-dependent (**Fig. 1D**). PDTOs cultured in Wnt-based medium often display a more cohesive, rounded and smooth organoid morphology (**Fig. 1E, bottom, Fig. S2C).** In contrast, organoids grown in OP medium show a more irregular, crenulated and less spherical shape and higher structural heterogeneity (**Fig. 1E, top, Fig. S2C)**.[6, 37]..

Importantly, key lineage markers remained preserved across both media (**Fig. S2D**), confirming maintenance of tumor-of-origin identity. For example, PDAC organoids retained KRT7/8/18/19, S100P, IGF2BP3 and MUC1, while CRC organoids expressed CDX2 and VIL1 with expected KRT7- negativity (and loss of KRT20 in MSI CRC014 [38]). Only minor medium-dependent variations were observed despite morphological differences.

To assess whether OP and Wnt culture conditions preserved the malignant genotype, WES was performed on a subset of 9 PDTO lines. Germline fingerprinting confirmed that matched parental, OP- and Wnt-derived samples were genetically related and showed no evidence of sample mix-up or cross-contamination (**Fig. S3A**). For two CRC and three PDAC patients, matched parental FFPE tumor tissue was available, allowing direct comparison of the mutational profiles of the original tumor and the corresponding PDTO cultures. Overall, a strong concordance was observed between parental tumor samples and derived PDTO lines in both CRC (**Fig. 1F**) and PDAC (**Fig. 1G**), indicating that the major tumor-associated mutations were retained during PDTO establishment and culture. An exception was observed for PDAC154, where several mutations, including KRAS and TP53, were detected in the PDTO cultures but not in the matched parental FFPE sample. This discrepancy may reflect lower tumor purity in the parental tissue sample, spatial tumor heterogeneity, or technical limitations related to variant detection in FFPE material. Notably, CRC018 was the only PDTO line that was established de novo in both OP and Wnt-based medium. In this model, the parental tumor tissue and both medium-specific organoid cultures showed highly concordant mutational profiles, supporting robust retention of the genetic background of the original tumor across both culture conditions (**Fig. S4**). Sequencing of additional matched OP- and Wnt-cultured PDTO lines showed highly concordant mutational profiles across culture conditions, indicating that both media largely preserved the genetic background of the PDTO models (**Fig. S5**).

### OP and Wnt media induce distinct transcriptional states while preserving tumor identity

To assess global transcriptional differences between culture conditions, we performed bulk RNAseq on the same 11 matched PDTOs derived from PDAC, CRC, GEJC, HNSCC and CUP tumors, grown in Wnt- versus OP media. UMAP analysis demonstrated that samples clustered predominantly by patient-of-origin rather than by medium, indicating preservation of tumor-intrinsic identity across conditions (**Fig. 2A, S3B**). Across tumor types, only a minority of genes was differentially expressed (e.g. FDR < 0.05; logFC 1): 3.9% in CRC, 8.5% in PDAC and 5.3% in GEJC (**Fig S6A**), similar to previous reports by Paul et al. [10]. Differential expression analysis revealed media-associated expression shifts (**Table S6: DEGs,** FDR < 0.001; LogFC 0.2): PDTOs cultured in Wnt medium upregulated proliferation-associated genes (e.g., STMN1, CDCA7, NAP1L1) and canonical Wnt-responsive stem cell regulators (LGR5, LRIG1) [39–43]. In contrast, OP-cultured PDTOs showed higher expression of epithelial state and adhesion-associated genes, including CEACAM6, COL17A1, AMIGO2 and AHNAK2 [44–48](**Fig. 2B**). Despite these state differences, cancer stem cell (CSC)-associated transcriptional signatures were broadly preserved across conditions, with only modest increases in the Wnt medium (**Fig. 2C**).

**Figure 2:**
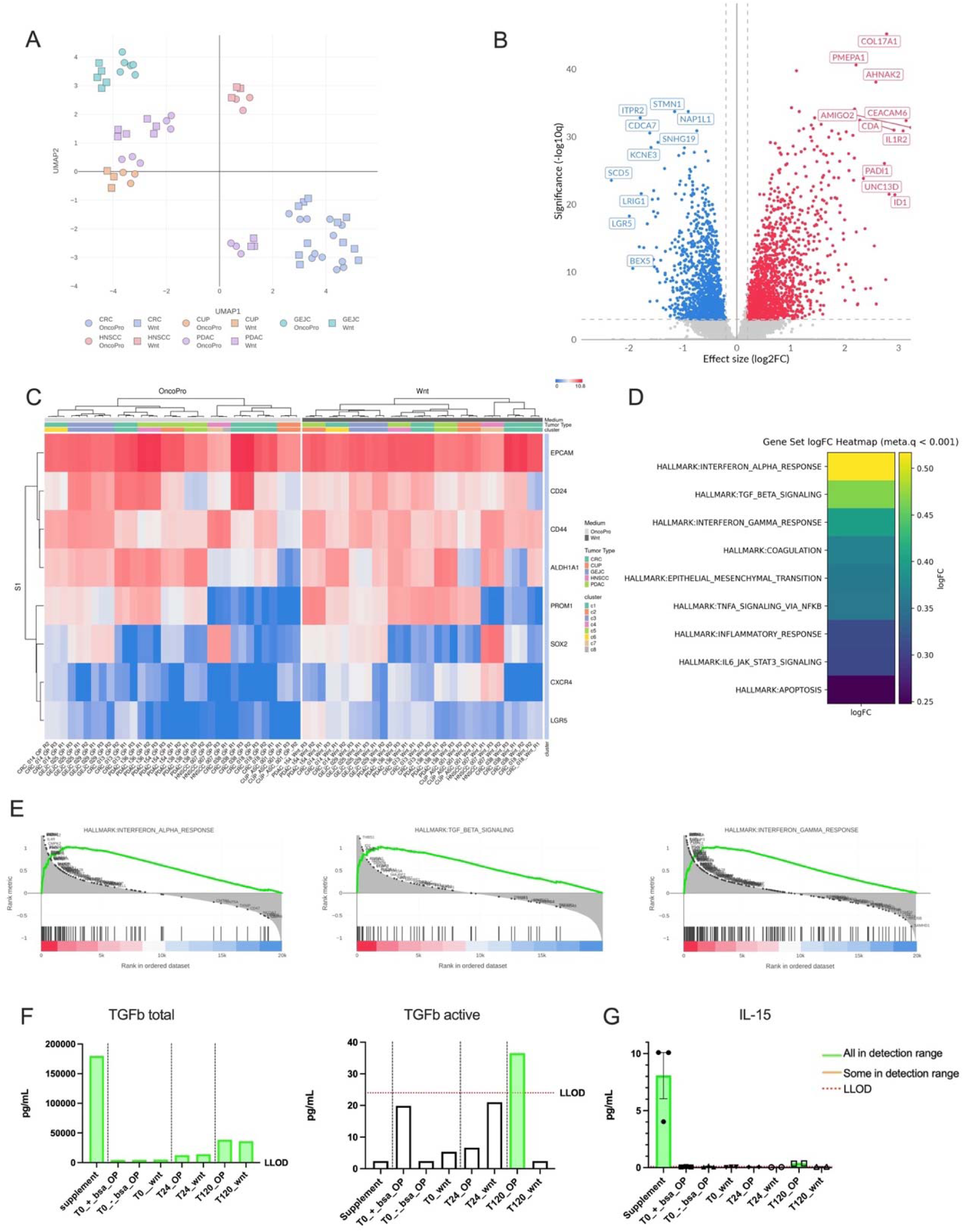
Culture medium drives distinct transcriptional states, with OP inducing inflammatory programs. (**A**) UMAP projection of bulk RNA-seq profiles from PDTOs, colored by tumor type (CRC (n=4), CUP (n=1), GEJC (n=2), HNSCC (n=1), PDAC (n=3)) and shaped by culture medium (circles = OP, squares = Wnt), showing that samples cluster primarily by patient of origin rather than medium. Each dot represents an individual RNA-seq replicate (three replicates per organoid line and medium condition); (**B**) Volcano plot of differentially expressed genes (DEGs) between OP and Wnt conditions. The x-axis represents effect size (log₂ fold-change OP vs Wnt) and the y-axis shows –log₁₀(q-value). Genes upregulated in OP (log₂FC > 0.2, FDR < 0.001) are highlighted in red; those upregulated in Wnt (log₂FC < –0.2, FDR < 0.001) are in blue. Selected top candidates are labelled. Analysis was performed on matched organoid pairs (n = 11 matched PDTO lines); (**C**) Heatmap of selected stemness-associated markers shows that both conditions maintain stem cell niche (n 1 matched PDTO lines). Absolute values, blue (downregulated); red (upregulated); (**D**) Heatmap of logFC values of Hallmark GSEA with metaq <0.0001 (n = 11 matched PDTO lines); (**E**) Hallmark GSEA demonstrating enrichment of inflammatory and cytokine-related signaling pathways in OP medium, including interferon-α, interferon-γ, and TGF-β signaling (n = 11 matched PDTO lines). All GSEA were significantly increased in OP (metq <0.001); (**F-G**) Mesoscale Discovery cytokine analysis of OP supplement, basal Wnt and OP medium (the latter with (+BSA) and without (-BSA) BSA)(T0), and supernatants from paired CRC018 PDTOs harvested at 24h and 120h. OP supplement contained high levels of total TGF-β. Active TGF-β increased during culture only in OP medium. IL-15 was present in the OP supplement. Values shown relative to assay lower limits of detection (red dotted line). <u>Abbreviations</u>: BSA, bovine serum albumin; CUP, carcinoma of unknown primary; CRC, colorectal cancer; GC, gastric cancer; GEJC, gastro-esophageal junction carcinoma; GSEA, gene set enrichment analysis; HNSCC, head and neck squamous cell carcinoma; LLOD, lower limit of detection; OP, OncoPro™ Tumoroid Culturing medium; PDAC, pancreatic ductal carcinoma; PDTO, patient-derived tumor organoid; SCC, squamous cell carcinoma; Wnt, Wnt-supplemented medium

Hallmark GSEA revealed significant enrichment of among others IFN-α response, IFN-γ response and TGF-β signaling in OP relative to Wnt-based culture conditions in the combined organoid cohort (FDR < 0.001; LogFC 0.2; metaq < 0.001) (**Fig. 2D–E, Table S7: GSEA**), suggesting increased cytokine- and TGF-β-related transcriptional activity in OP-cultured organoids. However, when applying stricter effect-size thresholds (e.g. logFC > 0.5, FDR < 0.05), only the IFN-α response remained significantly enriched, indicating that broadly shared medium-induced pathway-level differences across the full heterogeneous cohort are modest.

Further analysis of the genes contributing to the enriched pathways showed that their induction was patient- and tumor type-dependent (**Fig. S7, Table S7: GSEA**). For example, IFN-α-related genes were predominantly induced in OP-cultured PDAC organoids, whereas this pattern was not observed in CRC OP-cultured organoids (**Fig. S7**). These findings indicate that the global pathway enrichment reflects both recurrent medium-associated effects and tumor type-specific responses. Therefore, more focused tumor type-specific analyses were performed later for PDAC and CRC, for which sufficient sample numbers were available.

Given the cytokine-active state, we next performed an exploratory MSD assay comparing 49 cytokines (**Table S4: Cytokines**). Importantly, the OP supplement was first profiled directly, without prior contact with organoids, to determine which cytokines were introduced by the medium formulation itself. In addition, basal Wnt and OP media were analyzed at plating (T0), and supernatants from paired CRC018 PDTO cultures were collected after 24 h and 120 h (T24 and T120). This analysis showed that total TGF-β was already detectable in the OP supplement, indicating that TGF-β is directly introduced through the OP formulation rather than being solely produced by the organoids during culture. Active TGF-β was not detected at baseline but emerged during culture exclusively in OP supernatants, indicating activation of latent TGF-β during OP culture. No induction of active TGF-β was observed in Wnt supernatants, consistent with the absence of total TGF-β supplementation in the Wnt formulation (**Fig. 2F**). The OP supplement also contained detectable IL-15, although this was below the LLOD in basal OP medium (T0) (**Fig. 2G**). Several other cytokines and chemokines accumulated over time in culture, either in both media or preferentially in OP. For example, IP-10 (CXCL10) and IL-8 (CXCL8) levels increased over time in the supernatants, with substantially higher accumulation in OP than in Wnt medium, suggesting that PDTO epithelial cells actively secrete these chemokines under OP-induced conditions. However, because these measurements were not normalized for PDTO size or cellularity, these results should be regarded as exploratory and differential analysis should be interpreted with caution (**Fig. S8)**.

Together, these data indicate that OP and Wnt media do not alter tumor lineage identity, but instead impose distinct, transcriptional states characterized by differences in genes associated with proliferation, cytokine signaling and adhesion. Importantly, CSC-associated programs were broadly retained in both media, with modest enrichment of classical Wnt-driven stemness markers in Wnt-medium.

### OP-up genes preferentially map to malignant epithelial programs in patient single-cell atlases

To determine whether the medium-induced transcriptional states observed in bulk RNAseq reflect epithelial programs present in patient tumors, we benchmarked the OP- and Wnt-induced gene expression profiles against publicly available scRNAseq atlases of human PDAC and CRC patients. We performed differential expression analyses for three matched OP vs. Wnt PDAC organoid lines and four matched CRC organoid lines to verify whether medium-dependent transcriptional programs varied by tumor type. Overlap of the top 50 differentially expressed genes revealed minimal shared medium-responsive genes (Wnt n = 12; OP n = 14), indicating tumor-dependent transcriptional responses (**Fig. S6B-C**).

We first examined PDAC bulk RNA-seq expression of established classical and basal-like marker genes [49]. Marker genes of both subtypes were more pronounced in OP-cultured PDTOs relative to Wnt (**Fig. 3A-B**). This is in line with previous reports that show that classical and basal-like programs are not mutually exclusive and can co-occur within individual PDTOs [49]. We then derived OP-up and Wnt-up gene signatures from the PDAC bulk differential expression contrast (**Fig. 3C, Fig S6D**) and projected these signatures onto malignant, premalignant and non-malignant epithelial compartments in patient-derived scRNA-seq data. The OP-up signature mapped predominantly to malignant epithelial cells, with highest scores in classical cells and intermediate scores in basal-like cells and was largely absent in non-malignant acinar and ductal epithelium (**Fig. 3D**). In contrast, the Wnt-up signature showed minimal expression in malignant epithelial cells and instead aligned with non-malignant acinar/ductal populations (**Fig. 3E).** AUCell scoring of full 50-gene OP-up and Wnt-up modules confirmed these patterns at the single-cell level: OP-up scores were high and spatially concentrated within malignant epithelial regions (particularly in the classical subset), whereas Wnt-up scores were low in malignant epithelium and comparatively enriched in non-malignant epithelial neighborhoods (**Fig. 3F**). Taken together, these findings indicate that, in PDAC, the OP-associated transcriptional state closely matches malignant epithelial programs observed in patient tumors, whereas the Wnt-induced state aligned with programs enriched in non-malignant epithelial programs. Given the small number of PDAC PDO pairs, this analysis should be considered exploratory rather than broadly generalizable.

**Figure 3:**
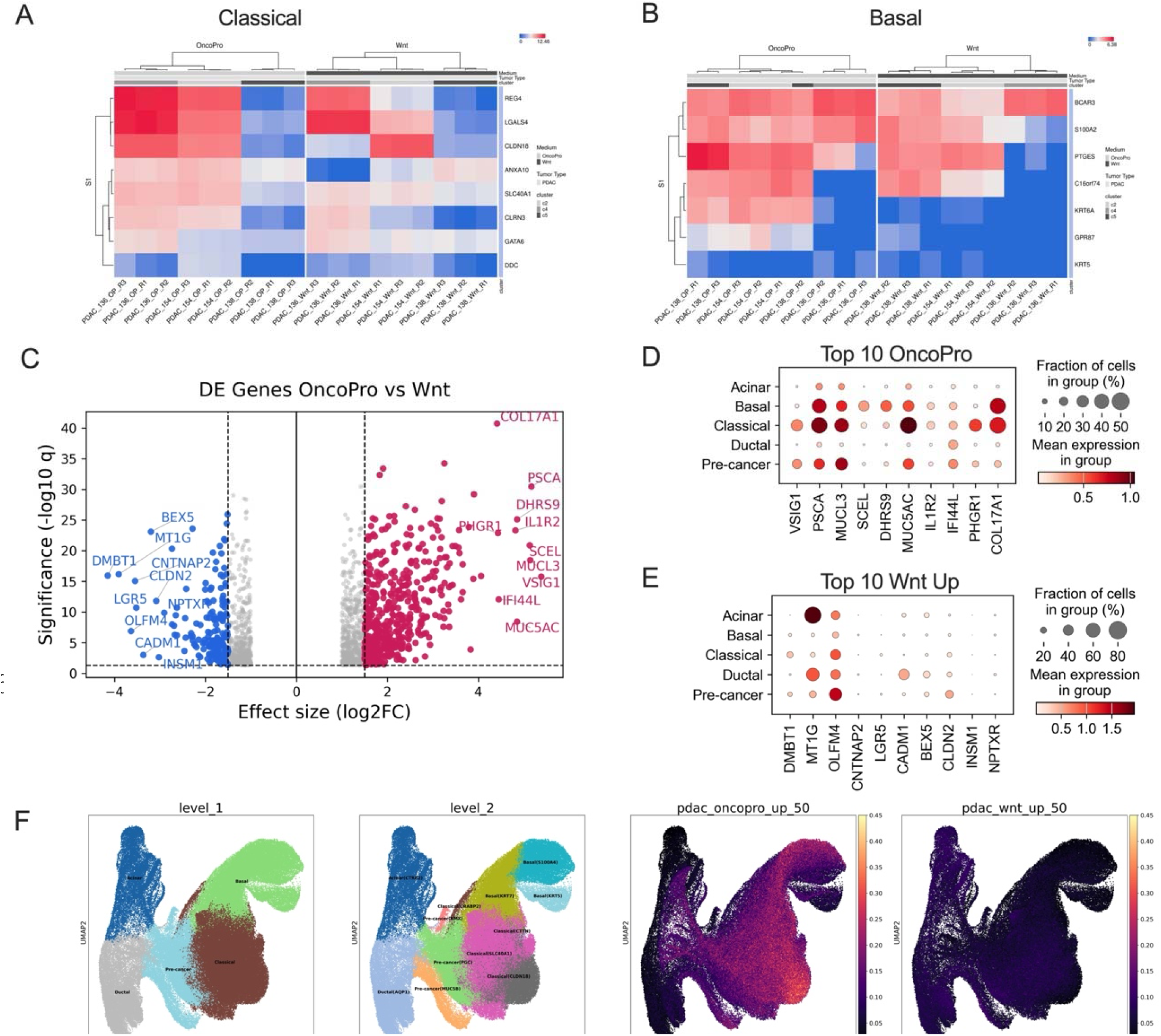
OncoPro*™* enriches for malignant epithelial state compared to Wnt in PDAC. (**A-B**) Heatmap of selected PDAC classical and basal like marker genes in paired PDAC PDTOs cultured in OP versus Wnt medium (n = 3 matched PDTO lines). Values expressed are absolute scores; (**C**) Differential expression analysis (volcano plot) of OP versus Wnt in 3 matched PDAC PDTOs. The x-axis displays effect size (log₂ fold-change), and the y-axis shows –log₁₀(q-value). Genes upregulated in OP (log₂FC > 1.5, FDR < 0.05) are indicated in red, and genes upregulated in Wnt (log₂FC < –1.5, FDR < 0.05) in blue. Top 10 upregulated genes per condition are annotated; (**D–E**) Projection of the top 10 OP-upregulated genes (D) and top 10 Wnt-upregulated genes (E) onto a reference single-cell atlas of human PDAC epithelium. Dot size indicates the fraction of cells expressing the gene in each epithelial state; color indicates mean expression. OP-induced genes preferentially localize to classical/basal malignant programs, whereas Wnt-induced genes align with acinar/ductal-like states; (**F**) Reference PDAC single-cell UMAP showing major malignant epithelial states (left), and AUCell scoring of OP-up and Wnt-up transcriptional signatures (right). OP signature scores are highest in classical/basal malignant compartments, while Wnt signature scores are enriched in non-malignant regions. <u>Abbreviations</u> DE, differentially expressed; FDR, false discovery rate; OP, OncoPro*Ô* Tumoroid Culturing medium; PDAC, pancreatic ductal adenocarcinoma; PDTO, patient-derived tumor organoid; scRNA-seq, single-cell RNA sequencing; UMAP, Uniform Manifold Approximation and Projection;

We next examined whether the medium-dependent transcriptional states observed in CRC organoids correspond to epithelial programs present in patient tumors. Differential expression analysis identified two gene sets: OP-cultured CRC PDTOs upregulated markers associated with epithelial state and EMT signaling (e.g., CEACAM5, CEACAM6, SERPINB5) [50–53], whereas Wnt-cultured organoids upregulated a smaller set of progenitor-associated markers (e.g., LGR5, LRIG1, SMOC2) [54–57] (**Fig. 4A**). To evaluate how these transcriptional states relate to patient tumor cell programs, we projected the top 10 OP-up and Wnt-up genes onto a large single-cell CRC atlas. The OP-up genes were expressed predominantly in malignant epithelial cells, with only slightly lower expression in healthy and polyp epithelium (**Fig. 4B**). Similarly, Wnt-up genes showed detectable expression in both malignant and non-malignant epithelial states (**Fig. 4C**). We next quantified the enrichment of the full 50-gene signatures using AUCell scoring across the epithelial compartment (**Fig. 4D; Fig. S6E**). The OP-up signature was primarily enriched in malignant tumor cells, whereas the Wnt-up signature remained low across malignant states, indicating that OP-associated signatures are more strongly enriched in CRC malignant cells in the analyzed atlas. Together, these findings indicate that CRC PDTOs cultured in OP medium are associated with a transcriptional state that aligns more closely with malignant CRC epithelial programs.

**Figure 4:**
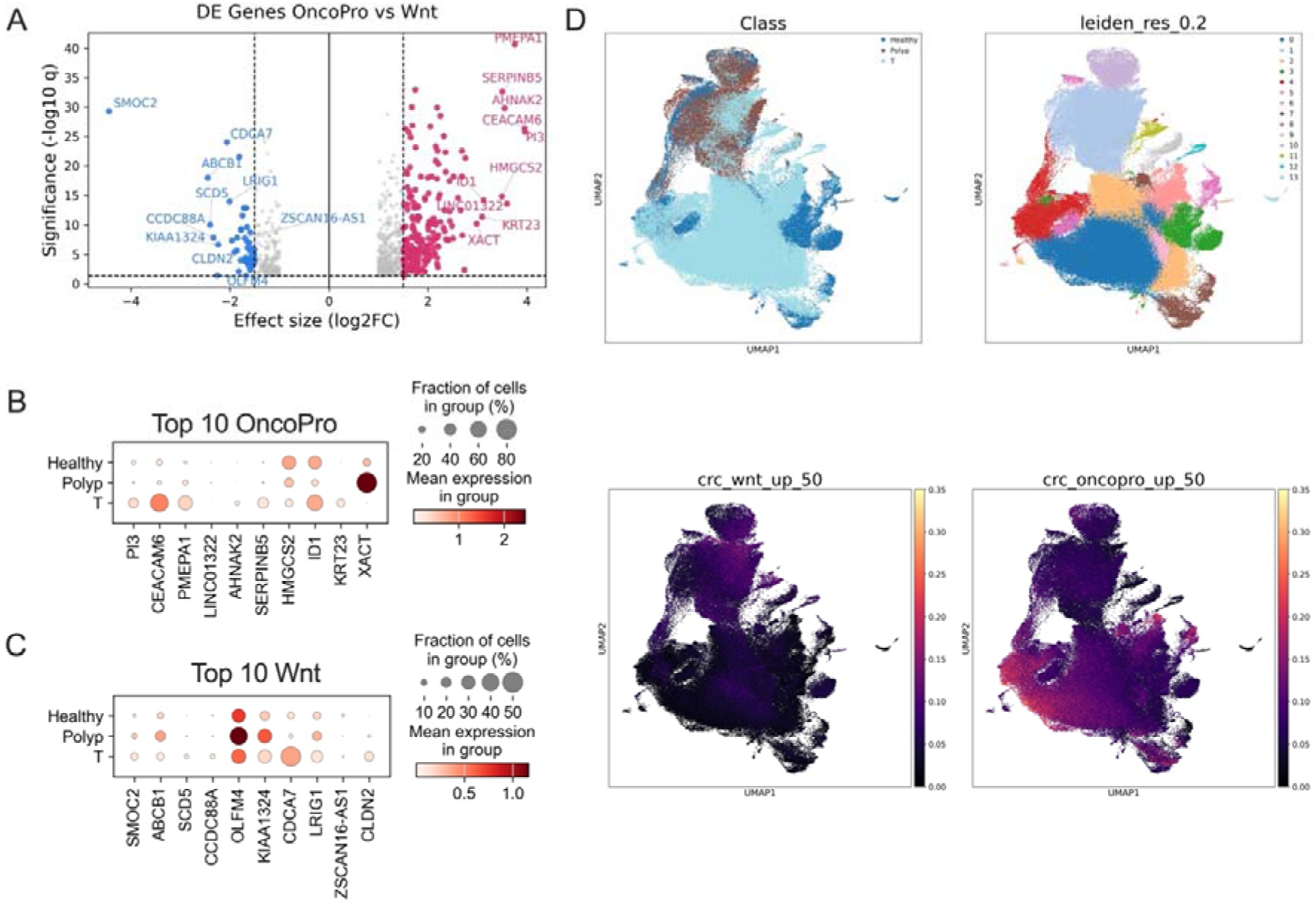
Medium-associated transcriptional programs in colorectal cancer organoids show differential representation across epithelial populations in a colorectal single-cell atlas. (**A**) Differential expression analysis (volcano plot) comparing CRC PDTOs cultured in OP versus Wnt medium (n = 4 matched PDTO lines). The x-axis shows log₂ fold change (OP vs Wnt), and the y-axis displays –log₁₀(q-value). Genes upregulated in OP are shown in red, genes upregulated in Wnt are shown in blue. Top 10 genes per condition are annotated; (**B–C**) Dot plots showing expression of the top 10 OP-upregulated genes (B) and top 10 Wnt-upregulated genes (C) across reference single-cell epithelial states from tumor, healthy colon and colorectal polyps. Dot size indicates the fraction of cells expressing each gene; color reflects mean expression; (**D**) Leiden clustering (top 50 genes per cluster Fig. S6E) and projection of OP-up and Wnt-up transcriptional signatures (AUCell scoring) onto a CRC scRNA-seq atlas annotated by malignant epithelial class an <u>Abbreviations</u>: CRC, colorectal carcinoma; DE, differentially expressed; FDR, false discovery rate; OP, OncoPro*™* Tumoroid Culture Medium; PDTO, patient-derived tumor organoid.

### Medium alters drug response in a patient- and class-dependent manner

We next investigated whether these medium-induced transcriptional states influenced drug response. Using transcriptomic signatures from the CTRPv2 and GDSC databases, we ranked 60 compounds by Normalized Enrichment Score (NES) and visualized media-dependent sensitivity predictions in a clustered heatmap (**Fig. S9A**). Two organoid clusters emerged: one (comprised exclusively of GEJC lines) with high NES variability and a second with more uniform profiles, in which OP-cultured PDTOs were predicted to be particularly sensitive to EGFR/HER2 inhibitors, Src inhibitors, statins and docetaxel, whereas Wnt-cultured PDTOs where thought to be more sensitive to PARP inhibitors, CDK4/6 inhibitors, Aurora kinase inhibitors, BCL2 inhibitors, irinotecan (SN-38) and cisplatin. To validate these in findings, we selected five matched PDTOs (PDAC138, PDAC154, CRC018, CRC038, CRC013) and performed an ex vivo drug screen using a panel that combined the predicted hit compounds with standard-of-care agents (**Fig. 5A**). Notably, two OP-cultured organoids (CRC013_OP and CRC038_OP) repeatedly expanded poorly, precluding their inclusion in high-throughput screening. Experiments were conducted on similar passage CRC and PDAC PDTOs, with larger passages differences in PDAC154 (**Table S3: passages**).

**Figure 5:**
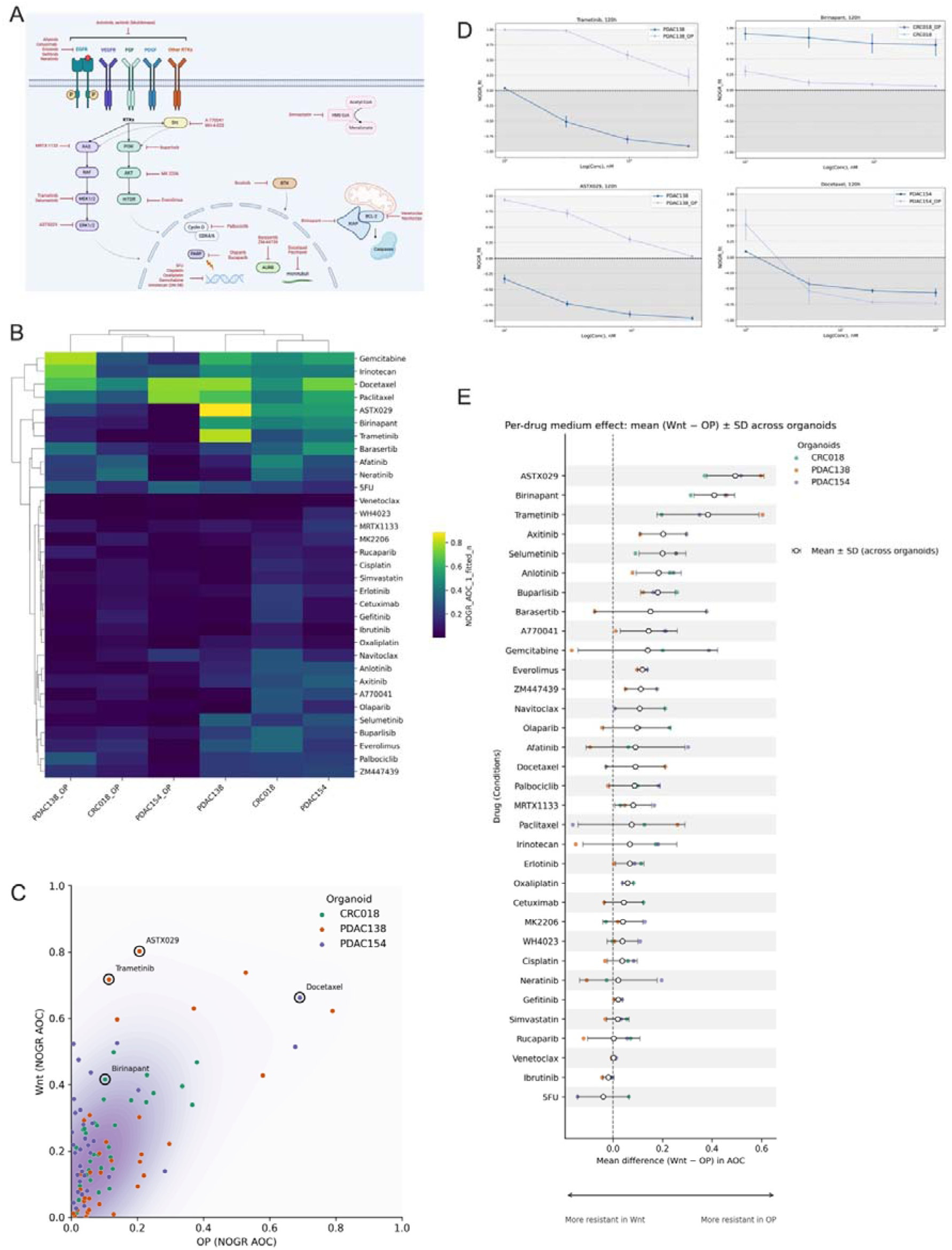
Media-dependent drug response in patient-derived tumor organoids (n = 3). (**A**) Schematic of major signaling modules and drug targets probed in the screen; (**B**) Clustered heatmap of drug responses. Hierarchical clustering of NOGR-AOC (0–1) across drugs (rows) and organoid-medium conditions (columns: CRC018, PDAC138, PDAC154 in OP and Wnt medium). Warmer colors indicate higher growth reduction. Clustering reveals medium-dependent sensitivities; Drug efficacy is represented by the normalized AOC ranging from 0, indicating no effect, to 1, indicating complete drug response (C) Scatter of mean NOGR-AOC for each organoid–drug pair (replicates averaged); OP on the x-axis, Wnt on the y-axis. A soft density background indicates point concentration. The four annotated points mark ASTX029, Trametinib, Birinapant and Docetaxel. Higher AOC values indicate greater drug sensitivity, whereas lower AOC values reflect relative resistance; (**D**) Exemplar NOGR dose–response plots, normalized to the matched vehicle and positive control, for ASTX029, trametinib, birinapant and docetaxel measured in matched OP and Wnt cultures across indicated organoids. Points show mean ± SD of replicates. Drug effects can be classified as: >1, proliferative effect; = 1, normal growth as in negative control; = 0, complete growth inhibition; = −1, complete killing as in positive control. Compounds were tested over log-scaled concentration ranges: 1–100 nM for docetaxel and 100–3000 nM for trametinib, ASTX029, and birinapant; (**E**) Per-drug medium effect (forest plot). For each compound, the open black circle denotes the mean ΔAOC = (Wnt − OP) across organoids; horizontal bars indicate SD (inter-patient variability). Small colored points are organoid-specific ΔAOC values. The dashed vertical line marks ΔAOC = 0. The bidirectional arrow below the x-axis indicates interpretation (*left*: more resistant in Wnt medium; *right*: more resistant in OP). <u>Abbreviations</u>: AOC, area over the curve; CRC, colorectal cancer; .NOGR, normalized organoid growth rate; OP, OncoPro™ Tumoroid Culturing Medium; PDAC, pancreatic ductal adenocarcinoma; PDTO, patient-derived tumor organoid; SD, standard deviation

Hierarchical clustering of the drug-response data in PDAC and CRC PDTOs (**Fig. 5B**) revealed segregation by culture medium rather than by individual organoid line, suggesting a substantial medium-associated effect on drug sensitivity. Across all paired organoid-drug measurements, OP and Wnt NOGR-AOC values were positively but only moderately correlated (Pearson r ≈ 0.59; p<0.001) (**Fig. S9B**). However, the joint distribution was skewed toward higher AOC in Wnt medium, pointing to a systematic medium effect with greater AOC values in Wnt medium, suggesting a generally more sensitive drug profile in Wnt medium (**Fig. 5C**). Representative dose-response curves along brightfield images for ASTX029, trametinib and birinapant demonstrated higher cell death in Wnt medium compared with OP across multiple PDTOs, whereas docetaxel showed minimal displacement (**Fig. 5D; Fig. S9D)**. In contrast, cisplatin and oxaliplatin showed essentially flat dose-response curves across the tested concentration range. Notably, these experimentally observed response patterns did not fully align with the initial sensitivity predictions from the CTRPv2 and GDSC transcriptomic signatures, possibly indicated that prediction models derived from large cell line-based datasets may not fully translate to organoid systems due to the inherent difference between both culturing systems. To quantify the medium effect per compound we summarized ΔAOC (Wnt – OP) across PDTOs (**Fig. 5E**). Several targeted agents showed consistently positive ΔAOC, most prominently MAPK-axis inhibitors (ASTX029, trametinib) and the IAP antagonist birinapant, indicating greater efficacy in Wnt medium. In contrast, taxanes (docetaxel, paclitaxel) and other classical chemotherapies (5-FU, irinotecan) centered near ΔAOC ≈ 0, suggesting relative medium insensitivity. Platinum agents (cisplatin, oxaliplatin) similarly showed minimal medium-dependent differences, although this reflected essentially flat dose-response curves across the tested concentration range, consistent with limited ex vivo activity in both media. The SD whiskers reveal clear inter-patient heterogeneity in the medium effect. In general, OP-grown PDTOs showed relative resistance compared to Wnt-grown PDTOs in the screened subset, especially in MAPK-inhibitors (**Fig. S9E)**.

To distinguish clonal selection from reversible state change, we performed two experiments: (i) We established a de novo Wnt line (PDAC138_WNT), transferred it to OP (PDAC138_WNT_OP) and back to WNT (PDAC138_WNT_OP_WNT) (**Fig. 6A**). (ii) We established two de novo organoid lines from the same parental sample: a Wnt line (CRC018_Wnt) and an OP line (CRC018_OP). The CRC018_OP line was subsequently transferred to Wnt medium (CRC018_OP_WNT) **(Fig. 6B**). All lines were then tested using the same drug panel to assess whether the sensitizing Wnt profile is induced by Wnt medium, or whether it is an intrinsic characteristic of the de novo Wnt-established lines.

**Figure 6:**
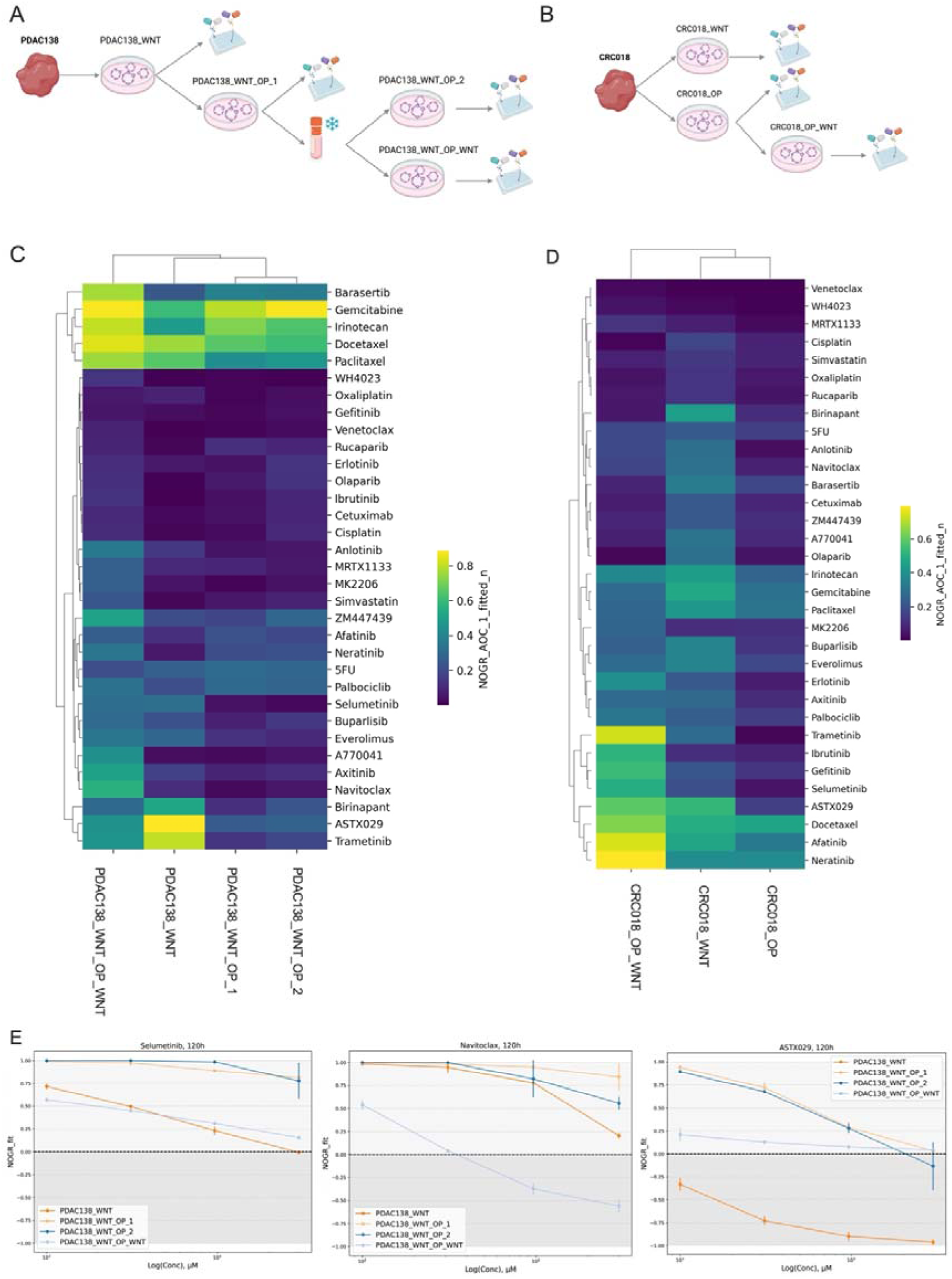
Functional plasticity of epithelial states underlies medium-dependent drug sensitivity in PDTOs (n = 2) (**A-B**) Schematic overview of medium transfer experiments. **(A)** A de novo PDAC organoid line was established in Wnt medium (PDAC138_WNT), transferred to OP medium (PDAC138_WNT_OP_1), after freezing & thawing we had PDAC138_WNT_OP_2 which was subsequently switched back to Wnt medium (PDAC138_WNT_OP_WNT). (**B**) Two de novo organoid lines were generated from the same parental CRC sample: one established in Wnt medium (CRC018_WNT) and one in OP medium (CRC018_OP). The CRC018_OP line was subsequently transferred to Wnt medium, yielding CRC018_OP_WNT; (**C**) Clustered heatmap of drug responses. Hierarchical clustering of NOGR-AOC (0–1) across drugs (rows) and organoid-medium conditions of PDAC138 Warmer colors indicate higher growth reduction; Drug efficacy is represented by the normalized area over the curve (AOC), ranging from 0, indicating no effect, to 1, indicating complete drug response; (**D**) Similar clustered heatmap for CRC018; (**E**) Exemplar dose-response plots for selumetinib, navitoclax and ASTX029 in PDAC138 during medium transfer experiments. Drug effects can be classified as: >1, proliferative effect; = 1, normal growth as in negative control; = 0, complete growth inhibition; = −1, complete killing as in positive control. Compounds were tested in a 4 point-titration from 100-3000nM, log scale <u>Abbreviations</u>: AOC, area over the curve; CRC, colorectal cancer; NOGR, normalized organoid growth rate; OP, OncoPro™ Tumoroid Culturing Medium; PDAC, pancreatic ductal adenocarcinoma; PDTO, patient-derived tumor organoid.

Hierarchical clustering of NOGR-AOC profiles placed PDAC138_WNT_OP_WNT (**Fig. 6C**) and CRC018_OP_WNT (**Fig. 6D**) on a distinct branch, apart from the parental Wnt condition, indicating a separate, transfer-induced state rather than simple convergence to either parental condition. Drug line plots show functional shifts toward greater sensitivity with three recurring patterns: (i) a full shift to Wnt-like sensitivity, (ii) a shift that exceeds the reference Wnt condition and (iii) minimal change, maintaining an OP-like response pattern. While these patterns were consistently observed across independent organoids, the specific drugs exemplifying each pattern differed between models (**Fig. 6E, Fig. S10A**). On average, sensitivity increased after retransferring to Wnt medium across the panel, arguing against predominant clonal selection (i.e., irreversible loss of sensitive subpopulations) and favoring medium-induced epithelial shift as the principal mechanism, while not fully excluding minor selection effects. Notably, drug response profiles were highly concordant between two independent PDAC138_WNT_OP experiments performed after cryopreservation and at different passage numbers, indicating strong reproducibility (pearman ρ = 0.92, p < 0.001; **Fig. S10B)**

Collectively, these data establish that Wnt-based formulation supports a state highly sensitive to targeted inhibition, whereas OP medium induces a phenotype that is generally more resistant, especially to MAPK-pathway and apoptosis-targeting therapies. Importantly, this state is at least partially reversible, emphasizing that medium composition dynamically reprograms epithelial function.

## Discussion

PDTOs are increasingly implemented in translational and preclinical research and previous reports showed the significant impact of culturing medium on morphology, transcriptomic profile and drug sensitivity [58, 59]. However, the impact of the novel OP medium on PDTO phenotype and therapeutic behavior has not yet been investigated. Here, we show that two epithelial organoid media, a classical Wnt/R-spondin-based formulation and the novel OP medium, impose distinct epithelial cell states while preserving overall patient- and tumor-type transcriptional and genomic identity. Importantly, these shifts likely reflect dynamic epithelial plasticity rather than fixed lineage conversion and further single-cell-resolved analyses will be required to fully delineate underlying state transitions.

In this study, transfer from Wnt medium to OP medium was successful in 83% of PDTO lines, comparable to previously reported rates [10], whereas de novo organoid establishment from fresh tissue reached only 11.1%, below the 50% success observed in the same study. Similarly, PDTOs retained tumor-of-origin transcriptional identity and more than 90% of genes were not differentially expressed between culture conditions. Nevertheless, the two media were associated with distinct phenotypic and pharmacologic states. Wnt medium was associated with enhanced drug sensitivity to several agents in the screened subset, whereas OP medium was associated with adhesion remodeling- and TGF-β signaling-associated programs, corresponding to broad drug resistance in PDAC and CRC PDTOs [60–62], particularly to MAPK-axis and apoptosis-targeting therapies. This is consistent with Raghavan et al. [59], who showed that PDTOs cultured in minimal medium were more resistant then PDTOs cultured in standard Wnt-containing organoid medium.

Importantly, the reversibility of drug sensitivity after medium switching argues against the irreversible loss of sensitive subclones and instead supports a dynamic, medium-dependent tumor cell state. These medium-associated transcriptional and pharmacologic differences were accompanied by distinct morphological features, suggesting that OP and Wnt conditions impose different signaling-dependent epithelial states. Tumor organoids lack organized lineage architecture, and morphological heterogeneity instead reflects underlying cell state and signaling context rather than a linear differentiation gradient [63, 64]. Consistent with this, both our bulk and single-cell transcriptomic analyses do not support terminal differentiation under OP conditions. To contextualize these reversible epithelial states in relation to patient tumors, we compared the medium-associated programs with public scRNA-seq atlases. In PDAC, the OP-associated transcriptional state mapped to malignant epithelial compartments, particularly the classical subtype, whereas the Wnt-induced state aligned with non-malignant acinar and ductal epithelium. In CRC, the OP-associated state spatially overlapped with malignancy associated programs. Together with the reversibility of drug sensitivity observed upon medium switching, these findings indicate that OP culture is associated with reversible redistribution of epithelial cell states, rather than fixed differentiation.

A plausible mechanism underlying the OP-associated inflammatory and TGF-β-related programs is medium-dependent activation of latent TGF-β. MSD analysis showed that the OP supplement contains high levels of total TGF-β, indicating that TGF-β is present within the proprietary supplement formulation. This is consistent with the canonical storage of mature TGF-β in latent complexes bound to Latency-associated peptide and latent TGF-β–binding proteins within extracellular matrix or cell-surface reservoirs [65, 66]. In matrix-embedded organoid cultures, matrix remodeling, protease activity, oxidative stress, or integrin-mediated tension could therefore promote activation of this latent TGF-β pool through established protease-, integrin/ECM-dependent mechanisms [65, 67, 68]. The presence of high total TGF-β in the OP supplement and the emergence of active TGF-β during OP culture are compatible with this model, but do not identify the responsible mechanism. Functional validation, including perturbation of TGF-β signaling and candidate activation pathways, will be required to determine whether TGF-β activation directly drives the observed inflammatory transcriptional states. Except for total TGF-β and IL-15 detected in the OP supplement, cytokine findings should be considered exploratory because conditioned suprnatants were not normalized for organoid cellularity.

These supplements are also relevant for future co-culture studies. In cancer-associated fibroblast (CAF)-containing models, the presence of active TGF-β (or TGF-β receptor inhibition in Wnt medium through A-83-01) is expected to markedly influence fibroblast activation state, ECM remodeling and paracrine signaling dynamics [69, 70]. Likewise, the detection of IL-15 in the OP supplement is noteworthy, as IL-15 is a key cytokine for natural killer (NK) cell survival, proliferation and cytotoxic priming [71, 72]. This may be particularly relevant for studies implementing CAR-NK or NK-tumoroid co-culture assays, where OP medium could directly modulate effector cell fitness [10]. Indeed, previous work has demonstrated that OP-cultured PDTOs can be readily combined with NK-based cytotoxicity assays, showing dose- and effector-to-target-dependent tumor cell killing in both primary NK cells and NK cell lines [10, 73, 74]. However, these systems typically rely on cytokine-enriched conditions (e.g., IL-2 supplementation and mixed media environments) [10, 73], underscoring that NK activity is not solely tumor-driven but strongly influenced by exogenous factors. In this context, the presence of IL-15 in OP medium may introduce additional confounding variables. While IL-15 primarily acts on immune cells, it has also been reported to influence cancer stem cell differentiation and chemotherapy sensitivity in certain tumor types [75]. Importantly, our group recently demonstrated that IL-15-secreting CAR-NK cells have superior cytotoxic activity against PDAC microtumors, composed of PDTOs and cancer-associated fibroblasts, compared with matched CAR-NK cells lacking IL-15 expression [76]. As a result, observed NK-mediated killing in OP-cultured PDTOs may partially reflect cytokine-driven activation rather than tumor-specific immune recognition. Together, these findings indicate that while OP medium is compatible with immune co-culture workflows, careful interpretation and experimental control are required to disentangle medium-induced immune activation from tumor-intrinsic responses.

The observed discrepancy between experimentally measured drug responses in PDTOs and predictions derived from GDSC/CTRP transcriptomic signatures reflects fundamental differences between these model systems. GDSC/CTRP databases were constructed from immortalized cell lines [77, 78], that undergo clonal selection under 2D culture conditions, resulting in fixation of specific oncogenic dependencies and loss of subclonal heterogeneity [79], whereas PDTOs show 3D architecture, cellular heterogeneity and tumor genomic fidelity [1–5]. These structural differences result in distinct pathway dependencies and metabolic profiles that influence drug sensitivity [79–82]. The 3D ECM in PDTOs creates gradients of oxygen, nutrients and drug penetration that significantly attenuate therapeutic effects compared to two-dimensional cultures [83–85]. PDTOs also maintain diverse, dynamic cell populations including CSCs that shift in response to microenvironmental cues, unlike the fixed populations in cell lines [86]. In addition, cell lines are maintained in fetal bovine serum-containing media, whereas PDTOs are cultured in defined, niche-factor-enriched media containing specific growth factors and pathway modulators. These differences in exogenous signaling context may further reshape transcriptional states, pathway dependencies and drug sensitivities. Consequently, GDSC/CTRP signatures require validation and adaptation for organoid-specific prediction models.

Retrospective studies have suggested that PDTOs can recapitulate patient drug responses, particularly for standard chemotherapy agents [87–89]. A recent meta-analysis reported a positive predictive value of approximately 68% and a negative predictive value of 78% across studies, indicating that organoid-based drug screening can stratify responders from non-responders with reasonable accuracy [90]. Notably, nearly all studies contributing to these estimates evaluated standard-of-care cytotoxic chemotherapies. Su et al. [91] also claimed that there is strong evidence for organoid sensitivity testing of 5-fluorouracil, irinotecan and to a lesser extent, oxaliplatin and TAS-102 in CRC. Importantly, these chemotherapies are also the drug classes in our dataset that exhibit minimal medium dependency, with ΔAOC values approximating zero across Wnt- and OP-grown PDTOs. For platinum agents specifically, however, this should be interpreted cautiously, as cisplatin and oxaliplatin showed largely flat dose-response curves across the tested concentration range. This suggests that the clinical concordance may, in part, reflect the robustness of cytotoxic drug responses to culture-induced cell state. In contrast, the few studies that attempted to predict response to novel targeted therapies report weaker or inconsistent clinical correlations [91]. For example, the SENSOR trial did not demonstrate meaningful predictive capacity for targeted treatments in metastatic CRC with vistusertib and capivasertib. Our results could give some insights in these discrepancies. Targeted therapies, including MAPK-pathway inhibitors, apoptosis-sensitizers and certain receptor tyrosine kinase inhibitors, show strong medium-dependent sensitivity shifts. Therefore, historical organoid-guided trials may have inadvertently screened targeted therapies in culture conditions that do not reflect the tumor state present in the patient, leading to apparent prediction failure due to artificial hypersensitivity.

Collectively, these findings highlight that culture medium is not a passive background but an active modulator of tumor cell state, drug response and clinical representativeness. Harmonization of media composition and transparent reporting of cytokine and growth-factor content could mitigate inter-laboratory variability [92]. Rather than advocating a single optimal medium, our data suggest that medium choice should be guided by experimental intent: Wnt-based media may be suited for expansion and discovery of stemness-associated vulnerabilities, whereas OP medium may capture malignant epithelial programs represented in patient single-cell atlases, which may be relevant for targeted therapy profiling. In settings of uncertainty, side-by-side testing across media systems remains warranted.

This study has several limitations. First, although the cohort included a wide range of tumor types, however the sample size within each subtype was modest, limiting statistical power for lineage-specific comparisons. In addition, several subgroup observations (e.g., MSI tumors, SCC, drug screenings) are based on limited sample numbers and should therefore be interpreted as descriptive rather than broadly generalizable. Second, cytokine profiling was restricted to a defined panel and only one CRC organoid line and should therefore be considered exploratory and hypothesis-generating. Additionally, because the OP medium is supplied as a proprietary formulation, its exact biochemical composition is not fully disclosed, which restricts mechanistic resolution of the pathways driving the observed state changes. Moreover, our comparative analysis focused primarily on differences between Wnt- and OP-grown PDTOs and did not extensively evaluate long-term phenotypic stability across passages. This aspect has been comprehensively addressed by Paul et al. [10], who demonstrated preservation of patient-specific molecular features and stability over time in OP-grown PDTOs. Next, only a subset of PDTOs were taken forward for in-depth morphological, transcriptomic, genomic and pharmacologic characterization. Because these represent PDTOs that were successfully established and maintained robust growth, this selection may introduce bias toward biologically fitter cultures and may underrepresent more fragile, but clinically relevant phenotypes. Furthermore, as drug responses were assessed within clinically achievable concentration ranges [93, 94], sensitivity that emerges only at supra-therapeutic doses may not be detected. However, such responses are unlikely to be clinically actionable and should be interpreted with caution when reported in in vitro systems [95]. Additionally, bulk RNA-seq limits resolution of cellular composition and hierarchical epithelial states and future single-cell analyses will be required to more precisely distinguish differentiation from reversible plasticity. In addition, while we demonstrate reversibility upon medium switch, the long-term stability and epigenetic consequences of these state transitions remain to be determined. Furthermore, we did not perform a systematic comparison with other defined organoid media formulations (e.g., surrogate Wnt-based systems or commercially available platforms). As the PDTO media landscape continues to evolve, broader cross-platform benchmarking studies will be important to further contextualize the positioning of OP relative to alternative culture strategies.

Future work should include expansion of the cohort, both in number of PDTOs and across additional tumor lineages to improve statistical robustness and generalizability. Larger and more diverse drug screens will be necessary to determine whether the observed media-dependent sensitivities extend across molecular subgroups and treatment classes. Mechanistic studies are needed to define the pathways underpinning the sensitivity phenotypes in the Wnt medium. Additionally, comparison of both media with regards to multiple co-cultures (e.g. effect of OP TGF-β supplementation on CAFs) is still lacking and would be of great interest. Moreover, scRNAseq profiling will be required to determine whether OP-associated transcriptional shifts reflect true reversible plastic state transitions, or selective outgrowth of specific subpopulations. Finally, prospective pairwise testing of Wnt- versus OP-cultured PDTOs against clinical treatment responses will be required to determine which media conditions most accurately recapitulate *in vivo* therapy outcomes for specific therapeutic classes.

## Conclusion

Culture medium is an important, controllable driver of epithelial cell state in PDTOs, with direct consequences for pharmacologic readouts. Across tumor types, OP and Wnt media preserved tumor-of-origin identity and oncogenic drivers yet imposed distinct transcriptional programs. In PDAC & CRC, Wnt medium was characterized by a broader drug sensitivity, whereas OP induced inflammatory signaling programs and aligned with malignant programs. The OP medium coincided with relative resistance in PDAC and CRC, especially to MAPK-axis and apoptosis-targeting agents. These functional differences were reversible upon medium switching, supporting state plasticity rather than fixed clonal selection.

Prospective studies linking parallel OP and Wnt medium screens to patient outcomes, expanded co-culture systems (e.g., CAFs, NK cells) under each medium, and mechanistic dissection of sensitivity shifts are likely to increase the reliability and clinical utility of organoid-guided precision oncology. Recognizing and controlling medium-induced plasticity is therefore essential for robust translational inference.

## Abbreviations

3D: three-dimensional
5-FU: 5-Fluorouracil
AC: adenocarcinoma
Ad-DF+++: Advanced DMEM/F12 supplemented with 1% GlutaMAX, 1% HEPES, 1% penicillin/streptomycin
AOC: area over the curve
BSA: bovine serum albumin
CAF: cancer associated fibroblast
CCA: cholangiocarcinoma
CRC: colorectal carcinoma
CSC: cancer stem cell
CUP: carcinoma of unknown primary
DEG: differentially expressed gene
EBUS-FNA: endobronchial ultrasound fine needle aspiration
ECM: extracellular matrix
FDR: false discovery rate
FFPE: formalin-fixed parafine embeded
GEJC: gastro-esophageal junction carcinoma
GSEA: gene set enrichment analysis
H&E: hematoxylin and eosin
HNSCC: head-and-neck squamous cell carcinoma
LLOD: lower limit of detection
MSD: mesoscale discovery
MSI: microsattelite instabile
NES: normalized enrichment score
NOGR: normalized organoid growth rate
NSCLC: non-small cell lung cancer
O: OncoPro™ grown organoid
OP: OncoPro™ Tumoroid Culture medium
PBS: phosphate-buffered saline
PDAC: pancreatic ductal adenocarcinoma
PDTO: patient-derived tumor organoid
RNAseq: RNA sequencing
SBS: single base substitution
SCC: squamous cell carcinoma
scRNAseq: single cell RNA sequencing
T: parental tissue
TPM: transcripts per million
UZA: University Hospital Antwerp
UMAP: Uniform Manifold Approximation and Projection
VAF: variant allele fraction
W: Wnt grown organoid
WES: whole exome sequencing

## Supplementary figures legends

**Supplementary Figure S1.**
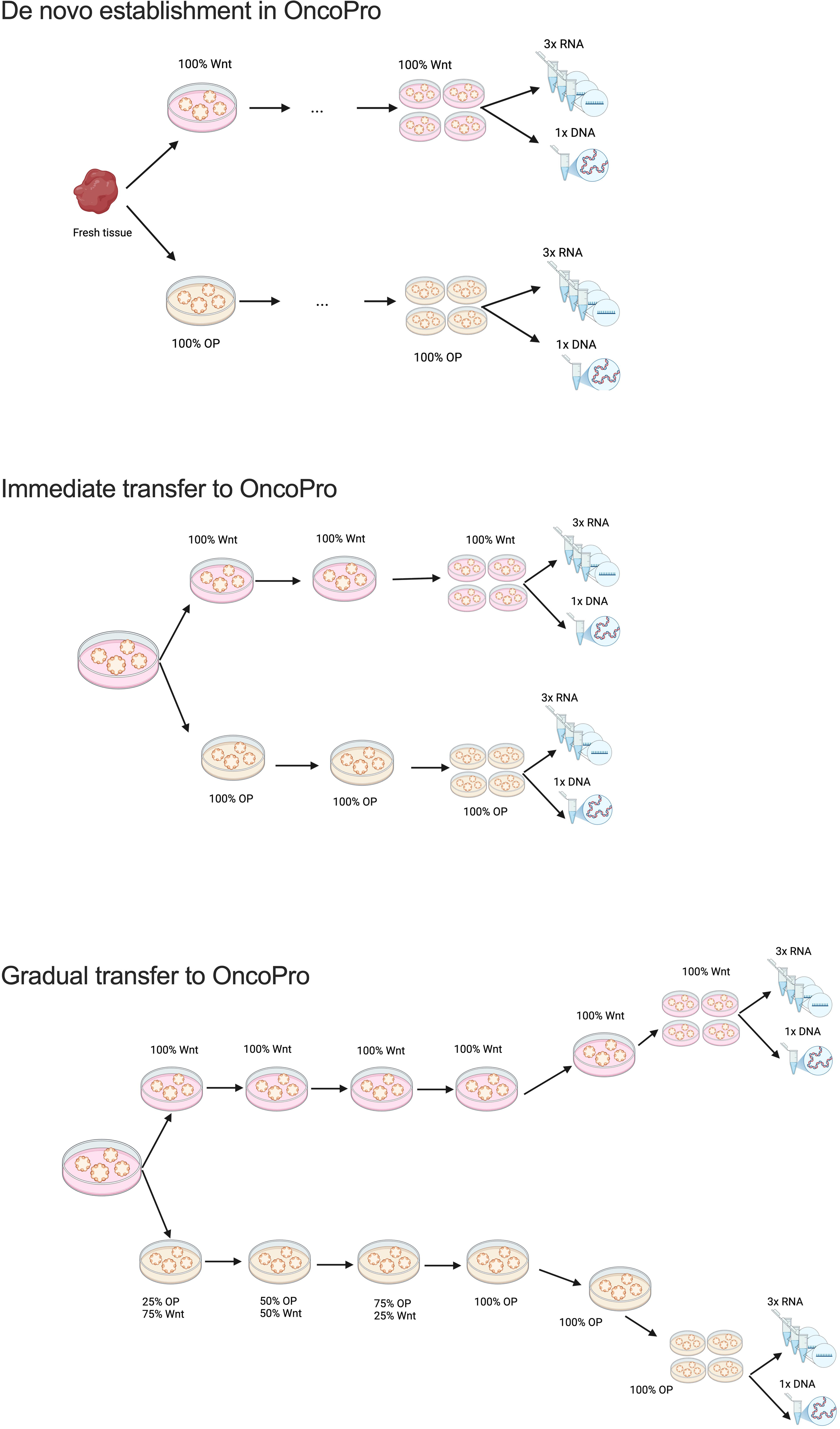
Schematic overview of PDTO establishment and transfer strategies in OncoPro and Wnt media. Schematic representation of three culture approaches: de novo establishment in Wnt or OncoPro (OP) medium, immediate transfer from Wnt to OP, and gradual adaptation to OP via stepwise medium transitions. For all conditions, paired PDTOs were maintained at comparable passage numbers prior to downstream RNA and DNA analyses <u>Abbreviations</u>: OP, OncoPro Tumoroid Culture Medium

**Supplementary Figure S2.**
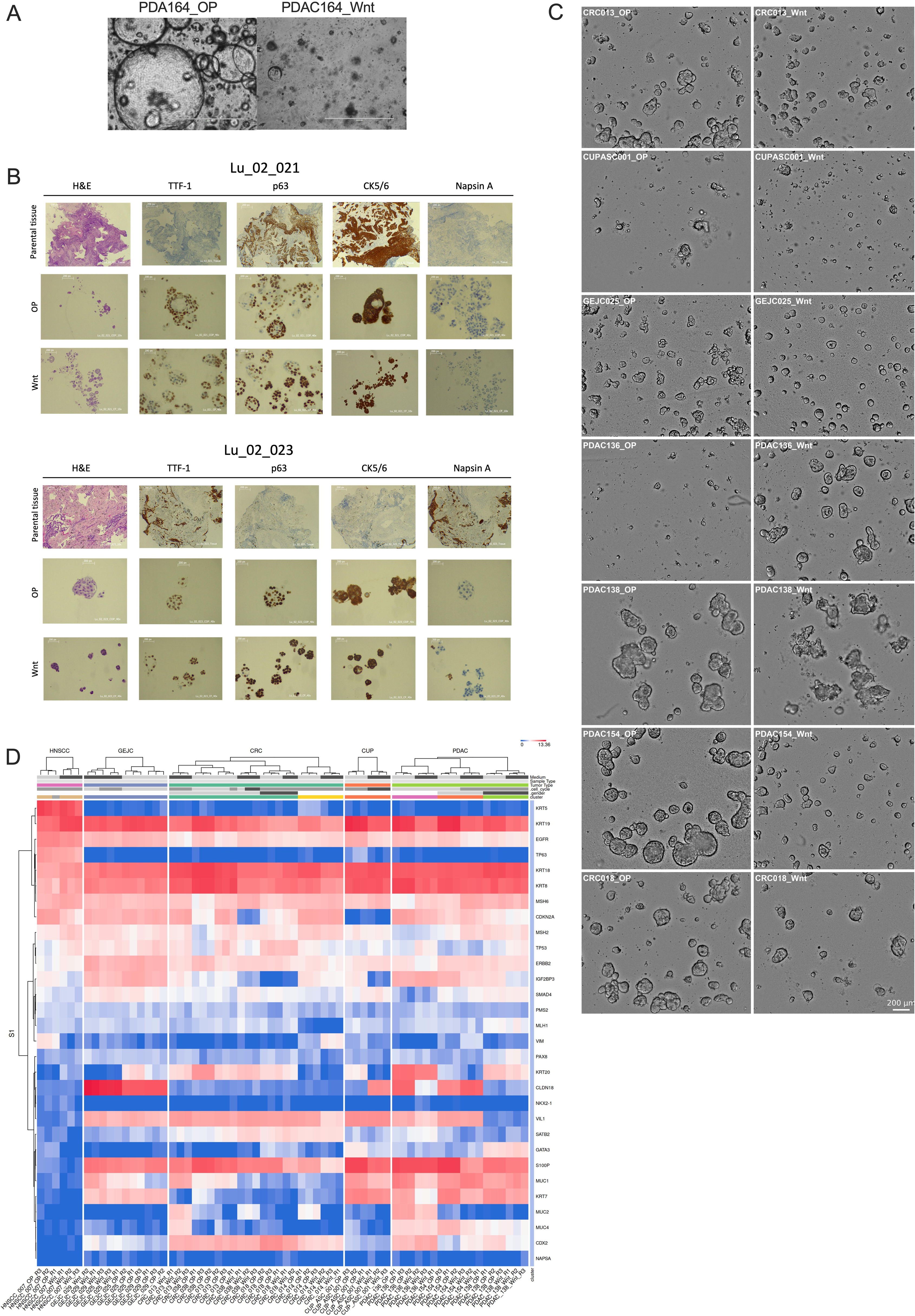
Phenotypes of PDTOs cultured in Wnt and OP media. A) Representative bright-field images captured shortly after organoid establishment. PDTOs cultured in OP medium exhibit larger and more cystic morphologies compared to those in Wnt medium; (B) Lineage marker immunohistochemistry (IHC) of two NSCLC carcinoma PDTOs (Lu_02_021 and Lu_02_023) cultured in OP and Wnt medium, compared with their parental tumor tissue. L_02_021 is a squamous carcinoma, while Lu_02_023 is an adenocarcinoma. Staining panels include hematoxylin and eosin (H&E), TTF-1, p63, CK5/6 and Napsin A, demonstrating non-malignant overgrowth; (C) Brightfield images of matched PDTO in Wnt and OP medium; (D) Expression heatmap of curated tumor lineage markers across PDTOs demonstrates retention of tumor-type–specific transcriptional identity independent of culture medium (absolute values; blue, downregulated; red upregulated). Relevant markers for PDAC (KRT7 (positive), KRT8/18/19 (positive), MUC1 (positive), S100P (positive)); CRC (CDX2 (positive), CK20 (positive, unless MSI), CK7 (negative or focal only), VIL1 (positive)); HNSCC (KRT5, TP63), GEJC (CK7, CLDN18), CUP (no clear markers). <u>Abbreviations</u>: CRC, colorectal carcinoma; CUP, carcinoma of unknown primary; GEJC, gastro-esophageal junction carcinoma; H&E: hematosin and Eosin-staining; HNSCC, head and neck squamous cell carcinoma; OP, OncoPro Tumoroid Culture Medium; PDAC, pancreatic ductal adenocarcinoma; PDTO, patient-derived tumor organoid

**Supplementary Figure S3.**
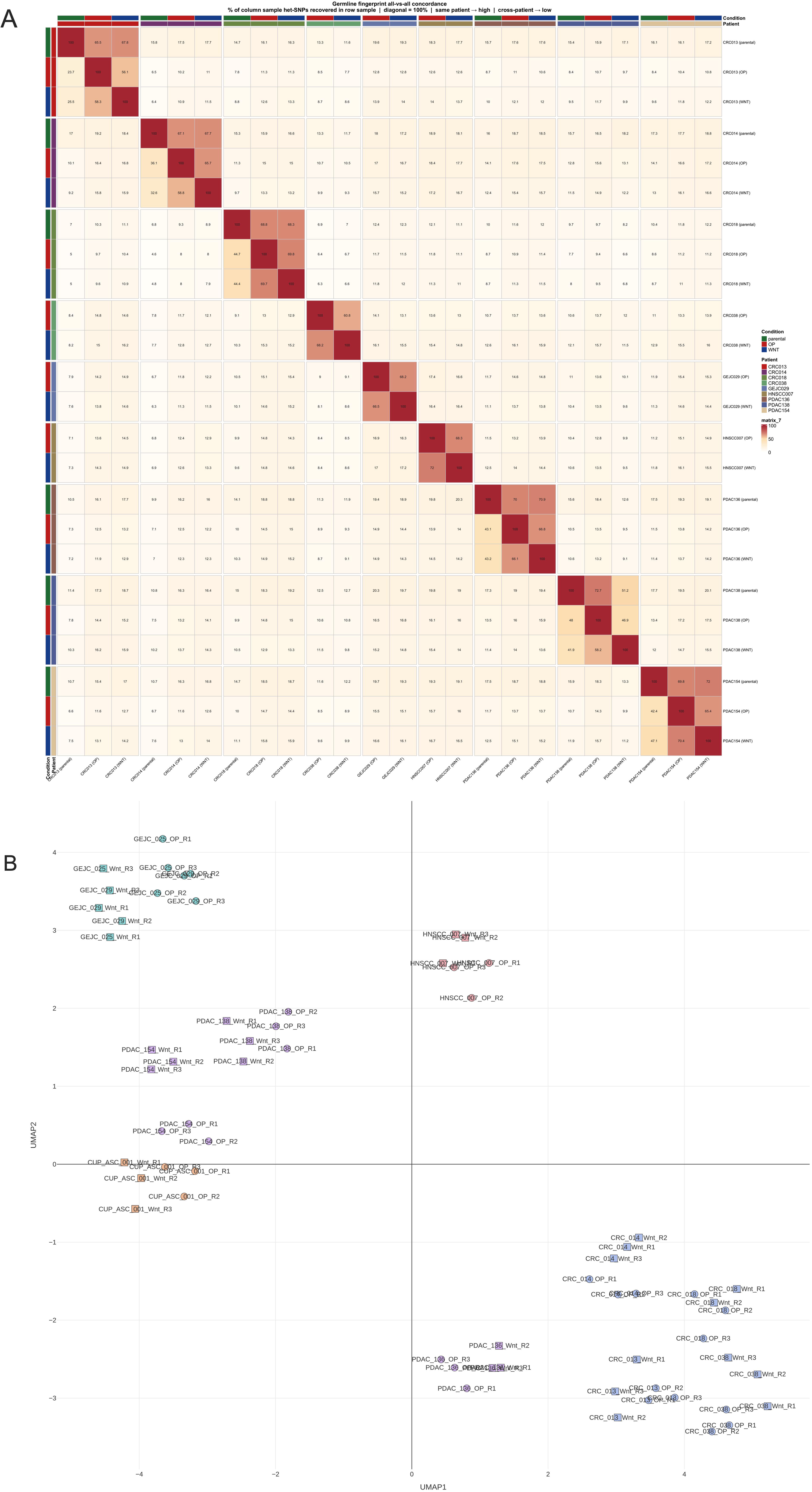
Genomic and transcriptomic profiling confirms patient-specific identity of matched Wnt- and OncoPro-cultured organoidMAP of bulk RNA-seq data with individual organoid line annotation. (A) Heatmap showing sample-to-sample genomic similarity based on germline fingerprint concordance cross patient-derived organoid cultures grown in Wnt-based or OncoPro medium. Each row and column represents one DNA-seq sample, and color intensity reflects the strength of the pairwise correlation, with darker red indicating higher transcriptomic similarity. Samples are ordered by patient/organoid line, revealing that matched samples from the same patient cluster together more strongly than samples from different patients. Annotation bars indicate culture condition and tumor type; (B) Uniform Manifold Approximation and Projection (UMAP) projection of bulk RNA-seq profiles from paired PDTOs cultured in Wnt and OncoPro (OP) media, with each point representing one replicate sample. Individual organoid lines are explicitly labeled to visualize sample-level relationships across conditions. Replicates cluster by organoid line, confirming matched-pair comparability across media, while medium-associated shifts remain secondary. <u>Abbreviation</u>: OP, OncoPro Tumoroid Culture Medium; UMAP, uniform manifold approximation and projection.

**Supplementary Figure S4:**
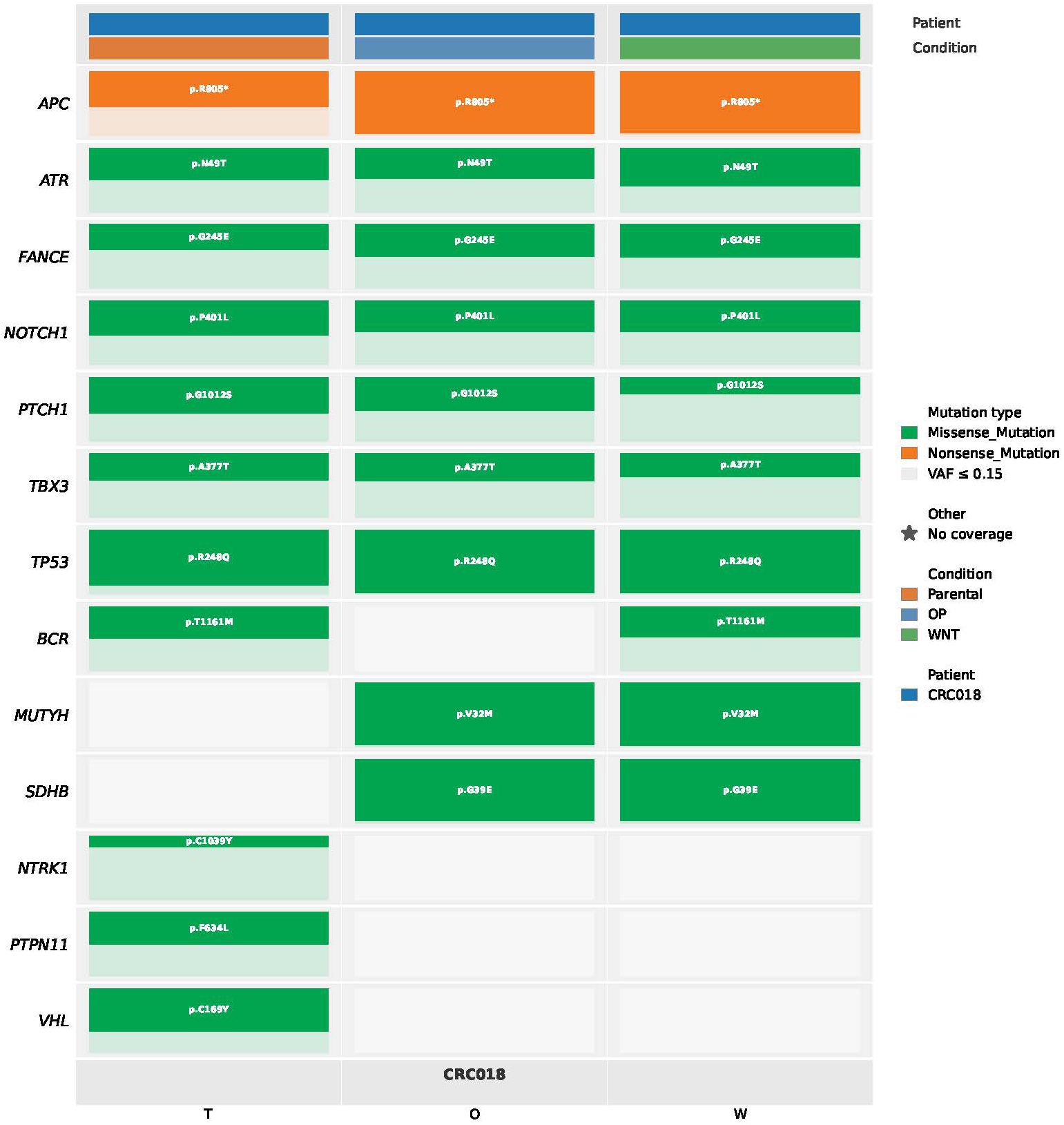
Shared variants support genomic concordance between parental tumor tissue and matched organoid cultures. Oncoplot showing variants detected in CRC018 parental tumor tissue and matched organoid cultures grown in OncoPro (OP) or Wnt-based medium (WNT). CRC018_OP and CRC018_WNT are both de novo derived. Columns represent the parental tumor and the corresponding organoid culture conditions, while rows indicate genes harboring detected variants. Colors indicate mutation type. Variant allele frequency (VAF) is indicated by shading, with low-VAF variants displayed separately, and missing coverage indicated where applicable. <u>Abbreviations</u>: CRC, colorectal cancer; OP, OncoPro; VAF, variant allele frequency.

**Supplementary Figure S5:**
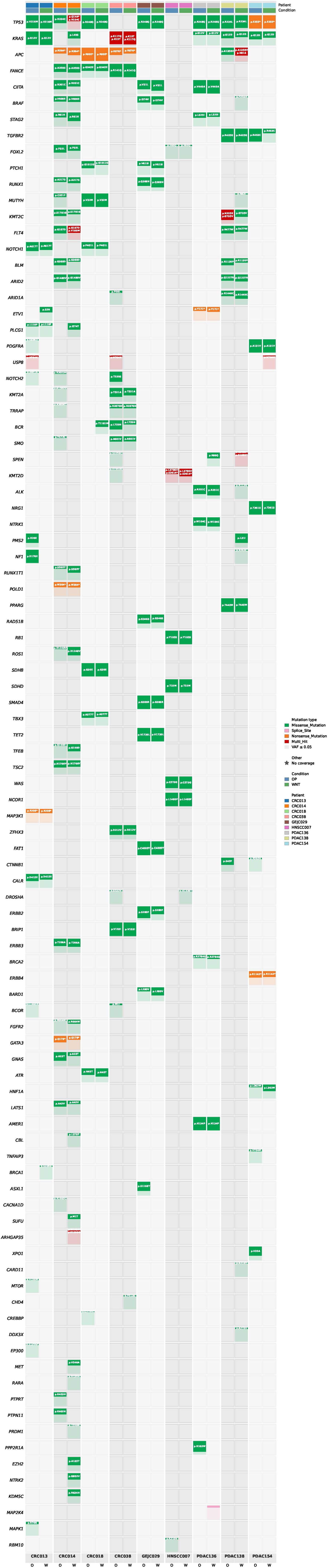
Paired OncoPro- and Wnt-cultured organoids retain patient-specific mutational profiles across tumor entities. Oncoplot showing variants detected in matched organoid cultures grown in OncoPro (OP) or Wnt-based medium (WNT) across nine patient-derived organoid lines. Columns represent paired culture conditions for each patient-derived line, and rows indicate genes harboring detected variants. Colors indicate mutation type. Variant allele frequency (VAF) is indicated by shading. The concordance of recurrent patient-specific alterations between matched OP and WNT cultures supports genomic stability and preservation of patient-of-origin identity across culture conditions. <u>Abbreviations</u>: CRC, colorectal cancer; OP, OncoPro; VAF, variant allele frequency.

**Supplementary Figure S6.**
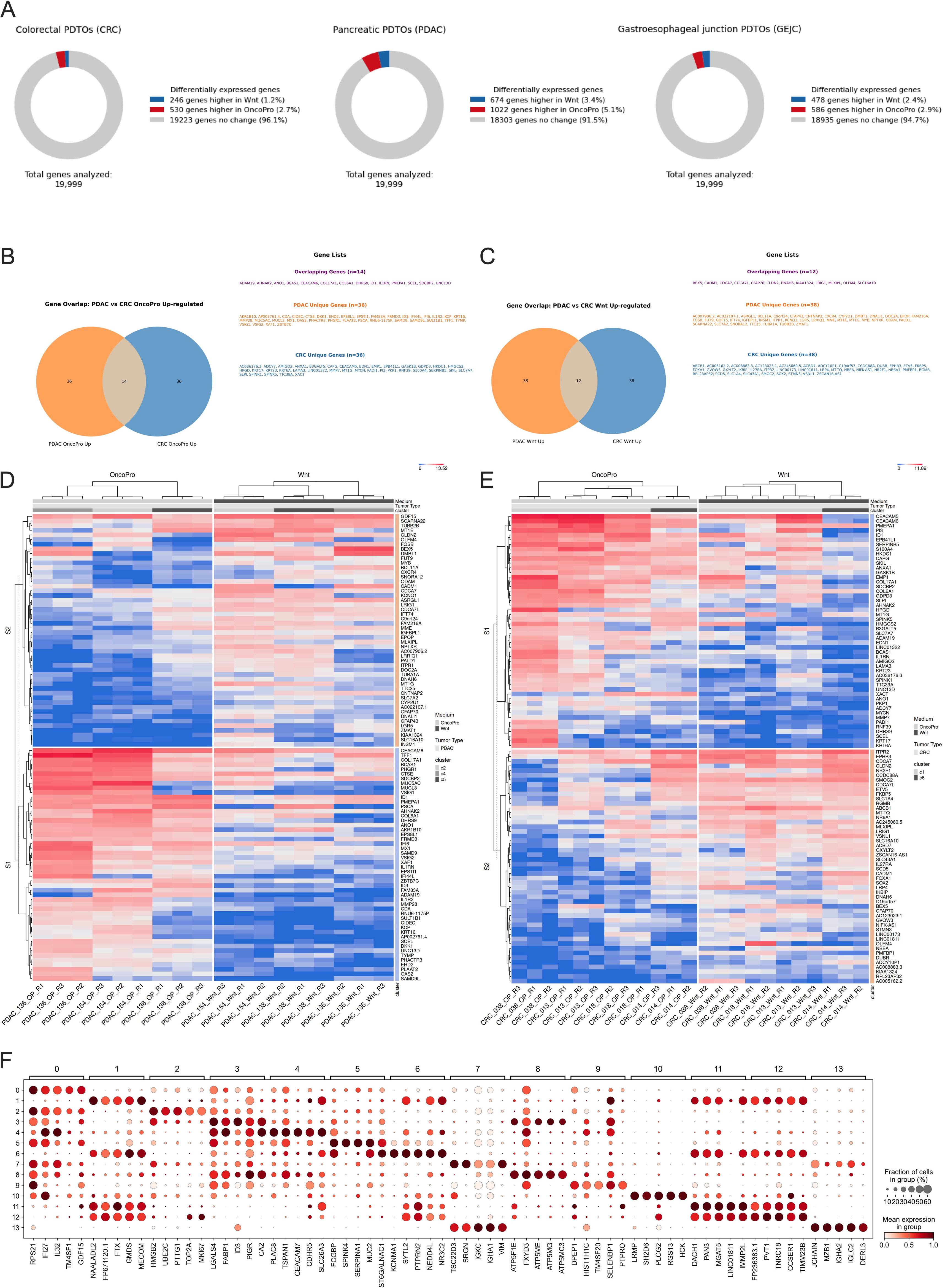
Medium- induced transcriptional states show tumor-type specific programs in PDAC and CRC PDTOs. (A) Donut plots showing the proportion of genes with higher expression in Wnt (blue) or OncoPro (red) medium, compared with non-differentially expressed genes (grey), across CRC, PDAC, and GEJC PDTOs. Percentages indicate the fraction of the total 19,999 genes meeting the differential expression thresholds (FDR < 0.05 and |logFC| > 1) (B-C) Venn diagrams showing the overlap in top 50 differentially expressed genes (DEGs) between PDAC and CRC PDTOs when comparing OP versus Wnt medium. Left: genes upregulated in OP; Right: genes upregulated in Wnt. Overlapping genes denote shared medium-associated transcriptional programs, whereas non-overlapping genes indicate tumor-type-specific responses. Representative examples of each category are listed; (D-E) Hierarchical clustered heatmaps of the Wnt_up_50 and OP_up_50 signatures PDAC (D) and CRC (E). Rows represent DEGs and columns represent individual PDTO samples, grouped by culture condition. Red indicates higher expression and blue indicates lower expression (absolute values); (F) Dot plot showing the expression patterns of key marker genes used to define the Leiden clustering across the CRC single-cell RNA-sequencing reference atlas **Abbreviations**: CRC, colorectal carcinoma; DEG, differentially expressed genes; PDAC, pancreatic ductal adenocarcinoma; PDTO, patient-derived tumor organoid; OP, OncoPro medium

**Supplementary Figure S7.**
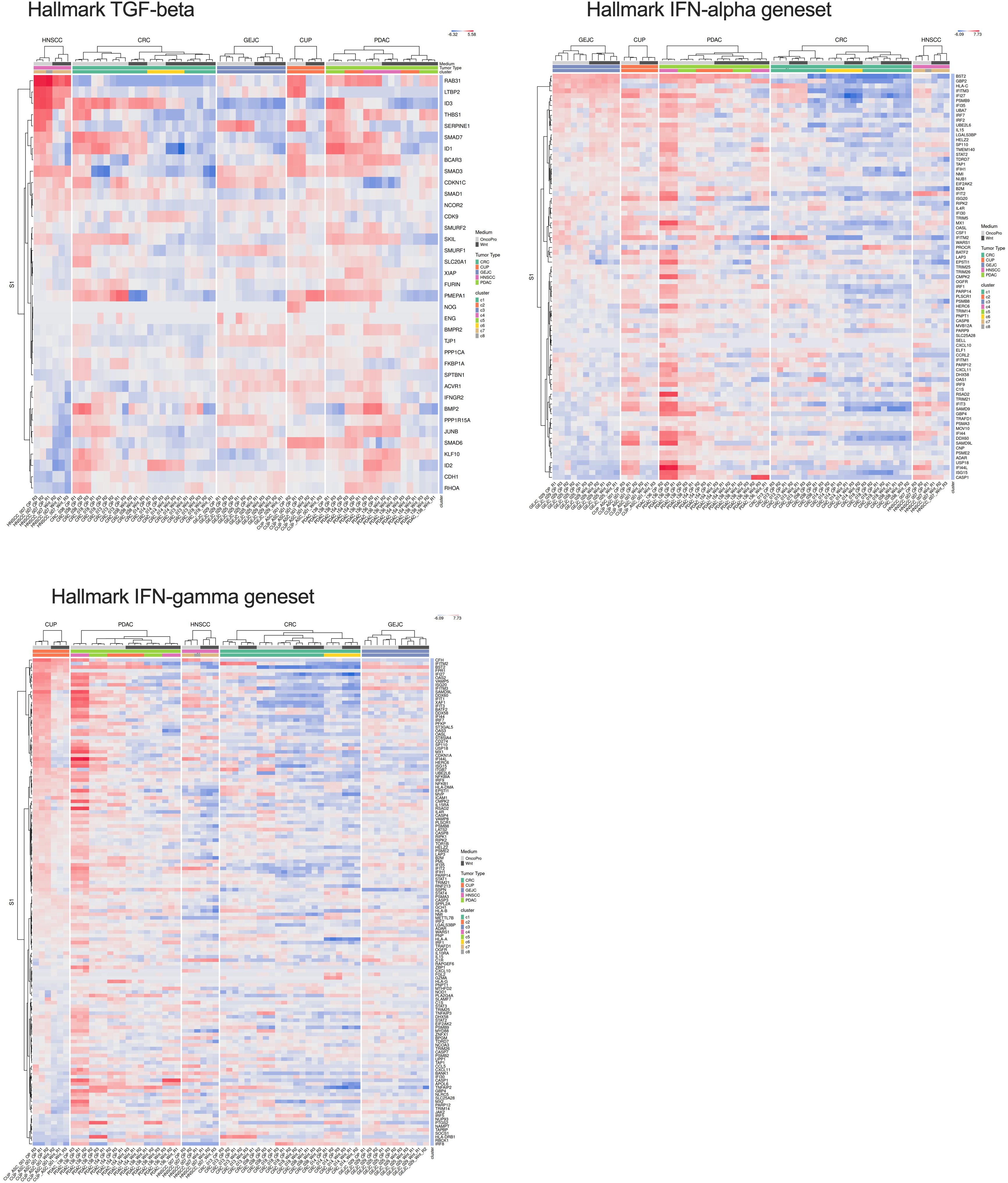
Patient- and tumor type-dependent expression of genes contributing to enriched inflammatory and TGF-β-related pathways. **Heatmaps showing the expression of genes included in the Hallmark TGF-**β, IFN-α and IFN-γ gene sets across PDTO samples cultured in Wnt-based or OP medium. Columns represent individual PDTO samples, annotated by tumor type and culture condition, and rows represent genes within the indicated Hallmark gene set.

**Supplementary Figure S8.**
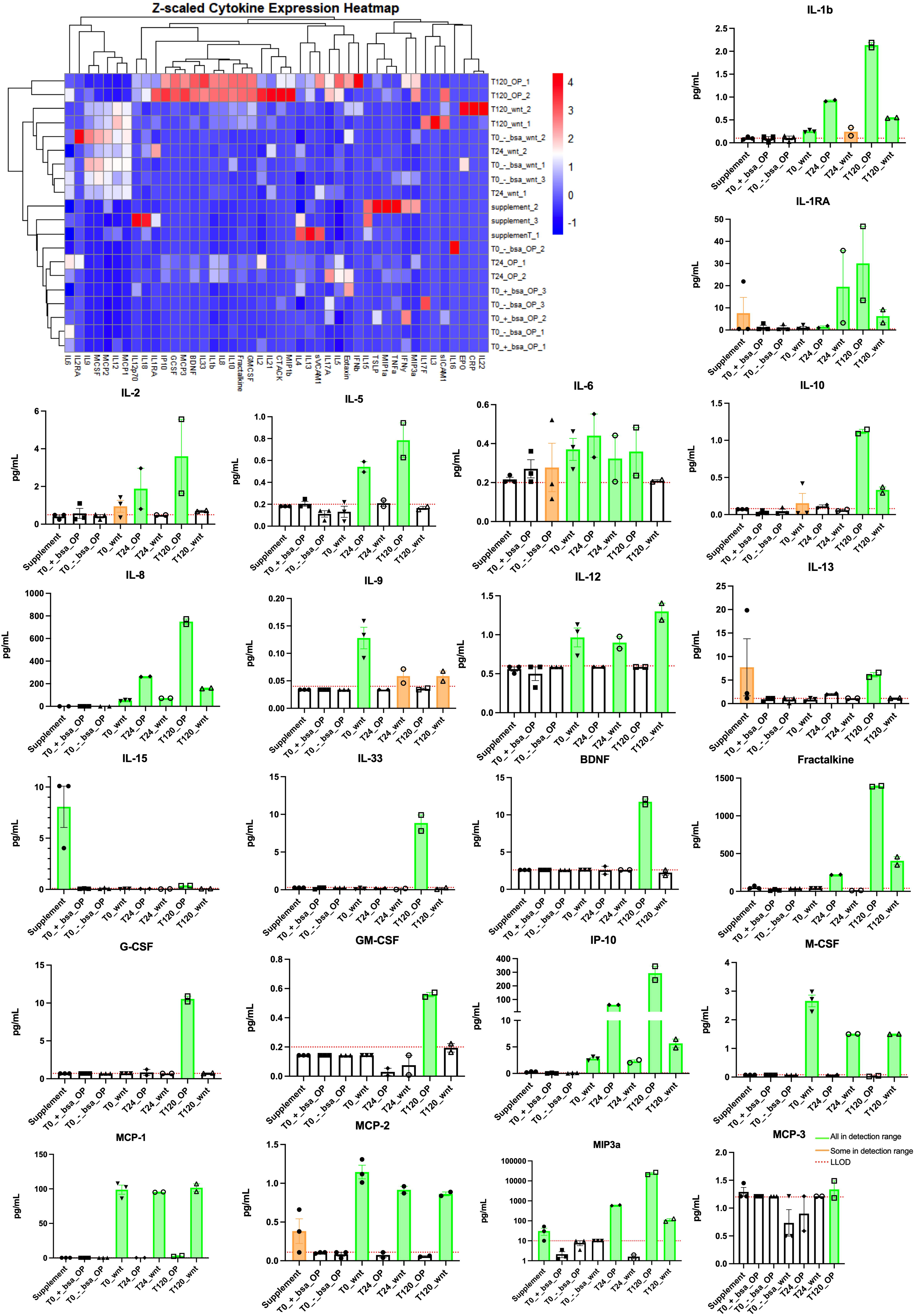
Mesoscale discovery of OP supplement, basal medium and supernatants. Hierarchically clustered Z-scaled heatmap of cytokine concentrations measured across unconditioned (baseline) and organoid-conditioned Wnt and OncoProÔ media at 0 h, 24 h and 120 h. A distinct OP T120h cluster was observed; Quantitative MSD cytokine profiles for selected representative factors. Bars represent mean cytokine concentration (pg/mL) ± SD across biological replicates; orange bars indicate values below the lower limit of detection (LLOD). CRC018 was used for this experiment. Not detected: IFNg, IL-4, IL-17A, CTACK, Eotaxin, IL12p70, TNFa, MIP-1a, EPO, IFNb, IL-12, IL-7, IL-16, IL-18, IL-17F; IL-21, IL-22, IL-3, TSLP, IL-2RA, MIP-1b, CRP, SAA, siCAM, sVCAM Abbreviations: BSA, bovine serum albumin; LLOD, lower limit of detection; OP, OncoPro Tumoroid Culturing Medium

**Supplementary Figure S9.**
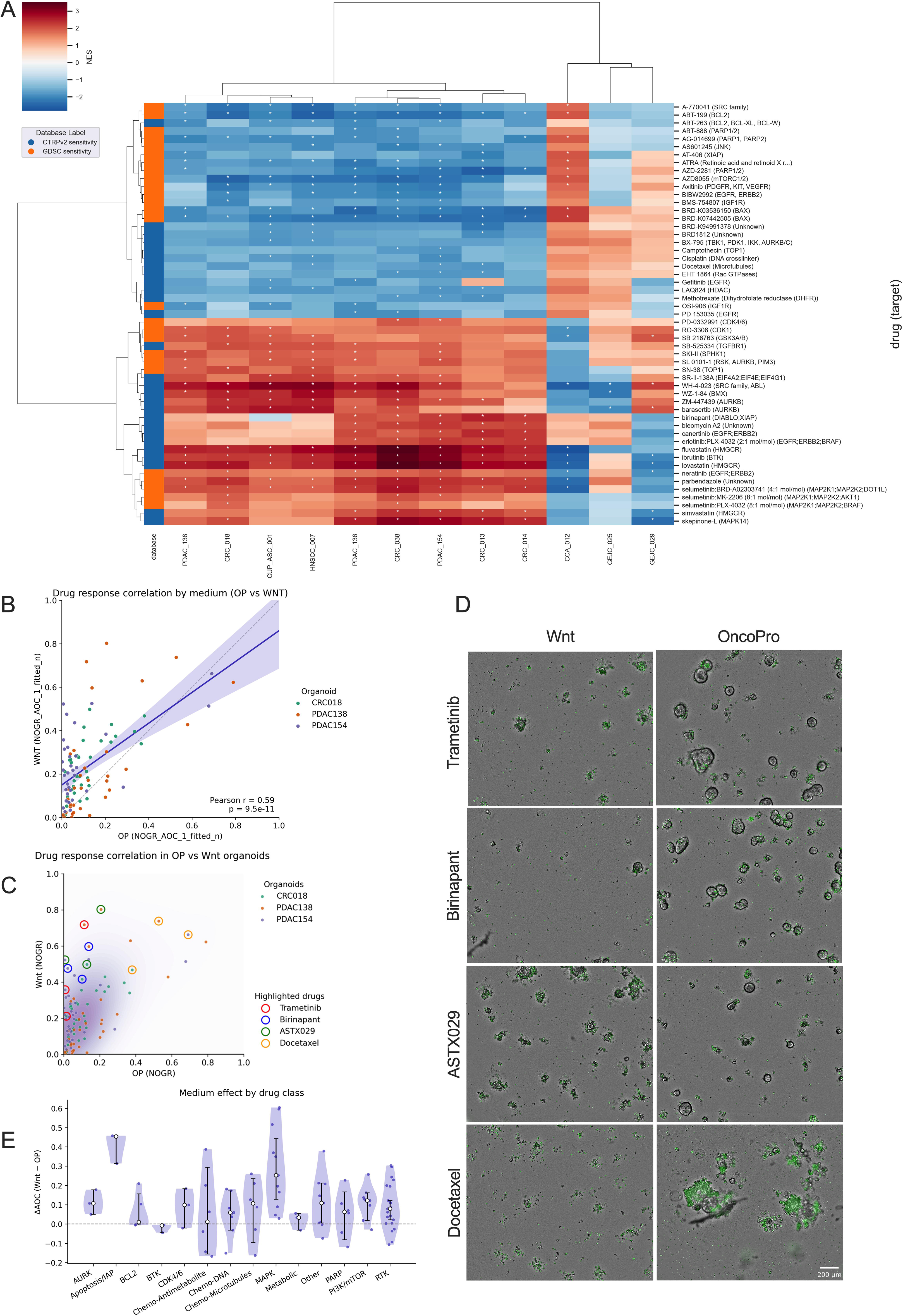
Drug screening on Wnt and OncoPro patient-derived organoids. (A) Clustered heatmap showing the predicted transcriptomic sensitivity profiles of 60 drugs across PDTOs. Drug response predictions were derived by comparing the measured gene expression profiles of PDTOs to drug-associated transcriptomic signatures from the CTRPv2 (blue) and GDSC (orange) databases. Drugs are grouped based on their Normalized Enrichment Scores (NES), with positive NES (red) and negative NES (blue) values represented by a gradient. Statistically significant results are marked with an asterisk. Rows represent drugs labeled with their targets, while columns represent PDTO lines grouped by hierarchical clustering; (B) Scatterplot of drug responses in OP- versus Wnt-grown organoids (mean NOGR_AOC_1_fitted_n values per drug). Each dot represents an organoid–drug pair, colored by organoid line. The blue line indicates the linear regression fit, with the shaded purple area representing the 95% confidence interval for the mean response. A moderate positive correlation was observed (Pearson r = 0.59, p<0.001). Drug efficacy is represented by the normalized area over the curve (AOC), ranging from 0, indicating no effect, to 1, indicating complete drug response; (C) same scatterplot as described above, only with density and important highlighted drugs. Trametinib (red), birinapant (blue) and ASTX029 (green), show increased sensitivity in WNT medium, while responses of docetaxel (orange) are generally well retained; (D) Brighfield images of the highlighted drugs in figure B; cytotox green stain shows cell death. Trametinib (PDAC138, 3000nM), birinapant (CRC019 3000nM), ASTX029 (PDAC138 3000nM), docetaxel (PDAC154, 100nM); (E) Violin plots show the per-class distribution; dots are individual organoid × drug observations; the white dot denotes the median with 95% bootstrap CI. The horizontal dashed line at 0 marks no medium effect. Abbreviations: AOC, area over the curve; CI, confidence interval; CRC, colorectal cancer; GEJC, gastro-esophageal junction carcinoma; HNSCC, head and neck squamous cell carcinoma; NES, normalized enrichment scores; NOGR, normalized organoid growth rate; OP, OncoPro Tumoroid Culture Medium; PDAC, pancreatic ductal adenocarinoma; PDTO, patient-derived tumor organoid

**Supplementary Figure S10.**
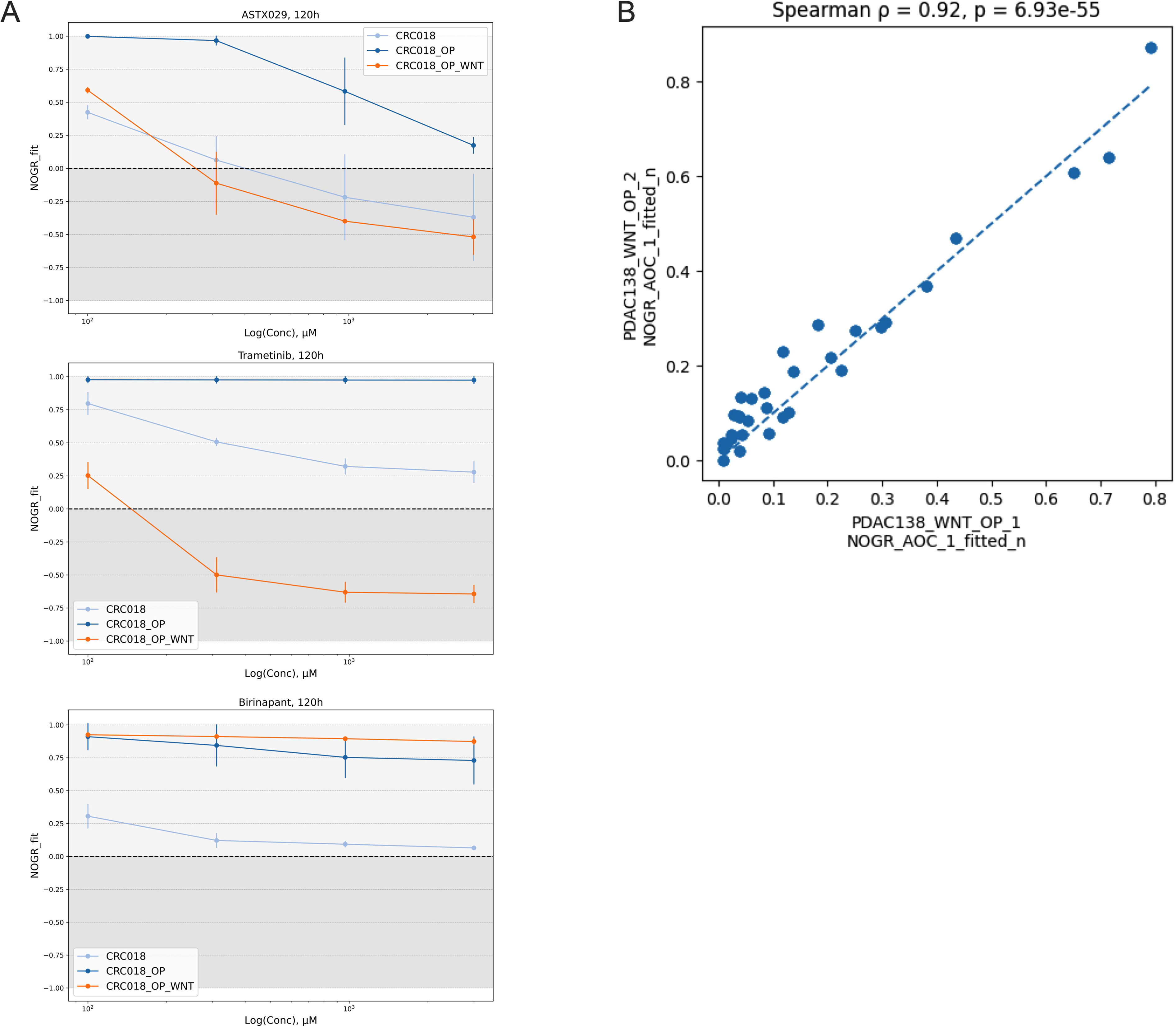
Functional plasticity. (A) Exemplar dose-response plots for selumetinib, navitoclax and ASTX029 in CRC018 during medium transfer experiments. (B) Correlation of drug response profiles between two independent PDAC138_WNT_OP experiments performed after cryopreservation and at different passage numbers. Each point represents a single compound, plotted as NOGR area-over-the-curve (AOC) values from the two experiments. A strong concordance was observed (Spearman ρ = 0.92, p <0.001), demonstrating high reproducibility of pharmacologic responses despite cryopreservation and passage differences. <u>Abbreviation:</u> AOC, area over the curve, NOGR, normalized organoid growth rate, OP, OncoPro Tumoroid Culture Medium;

## Acknowledgements

C.D. and S.S. conceived the study, designed the experiments and interpreted the results. S.S. and F.R. performed most experiments. S.S., F.R., M.L.C., J.B. and S.P. generated the PDTOs. G.R., V.H. S.V., W.T., N.K., G.V.H., J.H., P.V.S. and M.d.M provided human samples. S.K.S. and L.C. conducted MSD analysis. J.O. and C.D. conducted the whole exome sequencing analysis. S.S. wrote the manuscript. All authors reviewed and approved the manuscript.

## Data Availability

The data that support the findings of this study are available from the corresponding author, CD, upon reasonable request.

