## Supplemental Table 2 for "Comparative characterization of OncoPro™ and Wnt-Based media reveals distinct phenotypic and pharmacologic states in patient-derived tumor organoids"

Supplementary Table 2: Cell culture media

| **Medium** | **Components** |
| --- | --- |
| Standard medium | **Advanced DMEM/F12 supplemented with 1% GlutaMAX, 1% HEPES, 1% penicillin/streptomycin** |
| Collection medium | **Standard medium supplemented with 2% Primocin** |
| PDAC Wnt medium | Standard medium supplemented with **0.5 nM Wnt, 4% Noggin-conditioned medium, 4% Rspo-conditioned medium, 1x B27, 10 mM nicotinamide, 1,25 mM N-acetylcysteine, 100 ng/ml FGF-10, 500 nM A83-01,10 nM gastrin, and 10 μM Y-27632** |
| HN Wnt medium | **Standard medium supplemented with 4% Noggin-conditioned medium, 4% Rspo-conditioned medium, 1x B27, 10 mM, 1,25 mM N-acetylcysteine, 10 ng/ml FGF-10, 500 nM A83-01, 1 µM Forskolin, 50 ng/mL human EGF, 5 ng/mL FGF-2, 300 nM CHIR 9902, 1 µM Prostaglandin E2, 10 μM Y-27632** |
| GEJC Wnt medium | Standard medium supplemented with **0.5 nM Wnt, 4% Noggin-conditioned medium, 4% Rspo-conditioned medium, 1x B27, 10 mM nicotinamide, 1,25 mM N-acetylcysteine, 100 ng/ml FGF-10, 500 nM A83-01, 10 nM gastrin, 50 ng/mL hEGF, and 10 μM Y-27632** |
| CRC Wnt medium | Standard medium supplemented with **Noggin-conditioned medium, 4% Rspo-conditioned medium, 1x B27, 10 mM nicotinamide, 1,25 mM N-acetylcysteine, 100 ng/ml FGF-10, 500 nM A83-01, 10 nM gastrin, 50 ng/mL hEGF, and 10 μM Y-27632** |
| CUP Wnt medium | Standard medium supplemented with **Noggin-conditioned medium, 4% Rspo-conditioned medium, 1x B27, 10 mM nicotinamide, 1,25 mM N-acetylcysteine, 100 ng/ml FGF-10, 500 nM A83-01, 50 ng/mL hEGF, 1 μM SB202190, 25ng/ml FGF7 and 10 μM Y-27632** |
| Breast M1 Wnt medium | Standard medium supplemented with **Noggin-conditioned medium, 4% Rspo-conditioned medium, 1x B27, 10 mM nicotinamide, 1,25 mM N-acetylcysteine, 100 ng/ml FGF-10, 500 nM A83-01, 5 ng/mL hEGF, 1 μM SB202190, 5 ng/ml FGF7, 5nM hereguline b1 and 10 μM Y-27632** |
| Breast M2 Wnt medium | Standard medium supplemented with **0.5 nM Wnt Noggin-conditioned medium, 4% Rspo-conditioned medium, 1x B27, 10 mM nicotinamide, 1,25 mM N-acetylcysteine, 100 ng/ml FGF-10, 500 nM A83-01, 5 ng/mL hEGF, 10 nM forskolin, 100nM beta-estradiol, 500ng/mL hydrocortisone, 5nM hereguline b1 and 10 μM Y-27632** |
| Lung Wnt medium | **Standard medium supplemented with 4% Noggin-conditioned medium, 4% Rspo-conditioned medium, 1x B27, 10 mM nicotinamide, 1,25 mM N-acetylcysteine, 100 ng/ml FGF-10, 500 nM A83-01, 25 ng/mL FGF-7, 1 μM SB202190, 10 μM Y-27632** |
| Standard OP medium | OncoPro Basal medium supplemented with **1% penicillin/streptomycin** |
| PDAC/GEJC OP medium | Standard OP medium supplemented with 20 **μL/mL OncoPro Supplement, 62μL/mL OncoPro BSA, 10ng/mL OncoPro FGF-10,10 nM gastrin, 1x B27, and 10 μM Y-27632** |
| CRC OP medium | Standard OP medium supplemented with 20 **μL/mL OncoPro Supplement, 62μL/mL OncoPro BSA, 1x B27, and 10 μM Y-27632** |
| HNSCC/NSCLC/CUP OP medium | Standard OP medium supplemented with 20 **μL/mL OncoPro Supplement, 62μL/mL OncoPro BSA, 10ng/mL OncoPro FGF-10, 1x B27, and 10 μM Y-27632** |
| Breast OP medium | Standard OP medium supplemented with 20 **μL/mL OncoPro Supplement, 62μL/mL OncoPro BSA, 10ng/mL OncoPro FGF-10,10 nM beta-estradiol, 1x B27, and 10 μM Y-27632** |
| Digestion medium | Standard medium supplemented with **5 mg/mL collagenase II, and 10 μM Y-27632** |

Abbreviations: CRC, colorectal carcinoma; GI, gastro-intestinal; HN, head and neck; OP, OncoPro Tumoroid Culture medium; PDAC, pancreatic ductal adenocarcinoma
