## Supplemental Table 1 for "Comparative characterization of OncoPro™ and Wnt-Based media reveals distinct phenotypic and pharmacologic states in patient-derived tumor organoids"

### Supplementary Table 1: Reagents and resources

| **REAGENT and RESOURCE** | **SOURCE** | **IDENTIFIER** |
| --- | --- | --- |
| **Biological Samples** | | |
| Patient tumor pancreatic tissue | UZA | **14/47/480** |
| Patient tumor gastro-intestinal tumor | UZA | 2022-3470 |
| Patient tumor head and neck cancers | UZA | 20/08/090 |
| Patient tumor other cancers | UZA | 2023-5364 |
| **Peptides, recombinant proteins, and chemicals** | | |
| Advanced DMEM/F12 | Gibco | Cat#12634028 |
| GlutaMAX™ | Gibco | Cat#35050061 |
| HEPES | Gibco | Cat#15630080 |
| Penicillin-streptomycin | Gibco | Cat#15070063 |
| Primocin® | InvivoGen | Cat#ant-pm-05 |
| Collagenase type II | Sigma-Aldrich | Cat#C6885-1G |
| Y-27632 | Selleck Chemicals | Cat#S1049 |
| DPBS (10x) | Gibco | Cat#14200067 |
| Cultrex PathClear Reduced Growth Factor BME, Type 2 (10ml) | Tocris | Cat# 3533-010-02 |
| WNT surrogate-FcFusion protein | InvitroGen | Cat#PHG0402 |
| Noggin-Fc Fusion Protein conditioned medium | ImmunoPrecise Antibodies | Cat#N002-100ml |
| Rspo3-Fc Fusion Protein conditioned medium | ImmunoPrecise Antibodies | Cat#R001 - 100 ml |
| B27 Supplement | Gibco | Cat#17504044 |
| Nicotinamide | Sigma-Aldrich | Cat#N0636 |
| N-acetyl-L-cysteine | Sigma-Aldrich | Cat#A9165 |
| Human FGF-10 | Gibco | Cat#AF-100-26 |
| A83-01 | Tocris | Cat#2939 |
| Gastrin | Sigma-Aldrich | Cat#G9145 |
| Forskolin | Sigma-Aldrich | Cat#93049 |
| Human EGF | Stemcell | Cat#78136 |
| FGF-2 | Gibco | Cat#100-18B |
| CHIR 99021 | MedChemExpress | Cat# HY-10182 |
| Prostaglandin E2 | Tocris | Cat#2296 |
| OncoPro Basal Medium (part of OncoPro Tumoroid Culture Medium Kit) | Gibco | Cat#A5701201 |
| OncoPro Supplement (50x) (part of OncoPro Tumoroid Culture Medium Kit) | Gibco | Cat#A5701201 |
| OncoPro BSA (part of OncoPro Tumoroid Culture Medium Kit) | Gibco | Cat#A5701201 |
| FGF-10 Heat Stable | Gibco | Cat#PHG0372 |
| TrypLE™ Express | Gibco | Cat#12604-021 |
| Cultrex Harvesting Solution | Tocris | Cat#3700-100-01 |
| Recovery Cell Culture Freezing Medium | Gibco | Cat#12648010 |
| MycoAlert Mycoplasma Detection Kit | Lonza | Cat#LT07-518 |
| Tween 20 | Thermo Scientific Chemicals | Cat#233360010 |
| **Drugs** | | |
| Staurosporine | MedChemExpress | Cat#HY-15141 |
| 5FU | MedChemExpress | Cat#HY-90006 |
| A-770041 | MedChemExpress | Cat#HY-11011 |
| Afatinib | MedChemExpress | Cat#HY-10261 |
| Anlotinib | Selleck Chemicals | Cat#S8726 |
| ASTX029 | MedChemExpress | Cat#HY-126288 |
| Axitinib | MedChemExpress | Cat#HY-10065 |
| Barasertib | MedChemExpress | Cat#HY-10065 |
| Birinapant | MedChemExpress | Cat#HY-10065 |
| Buparlisib | MedChemExpress | Cat#HY-70063 |
| Cetuximab | MedChemExpress | Cat#HY-P9905 |
| Cisplatin | MedChemExpress | Cat#HY-17394 |
| Docetaxel | MedChemExpress | Cat#HY-B0011 |
| Erlotinib | MedChemExpress | Cat#HY-50896 |
| Everlolimus | MedChemExpress | Cat#HY-10218 |
| Gefitinib | MedChemExpress | Cat#HY-10218 |
| Gemcitabine | MedChemExpress | Cat#HY-10218 |
| Ibrutinib | MedChemExpress | Cat#HY-10997 |
| MK2206 | MedChemExpress | Cat#HY-10358 |
| MRTX-1133 | MedChemExpress | Cat#HY-134813 |
| Navitoclax | MedChemExpress | Cat#HY-10087 |
| Neratinib | MedChemExpress | Cat#HY-32721 |
| Olaparib | MedChemExpress | Cat#HY-10162 |
| Oxaliplatin | MedChemExpress | Cat#HY-17371 |
| Paclitaxel | MedChemExpress | Cat#HY-B0015 |
| Palbociclib | MedChemExpress | Cat#S4482 |
| Rucaparib | MedChemExpress | Cat#HY-10617A |
| Selumetinib | MedChemExpress | Cat#HY-50706 |
| Simvastatin | Selleck Chemicals | Cat#S1792 |
| SN-38 | MedChemExpress | Cat#HY-13704 |
| Trametinib | MedChemExpress | Cat#HY-10999 |
| Venetoclax | MedChemExpress | Cat#HY-15531 |
| WH-4-023 | MedChemExpress | Cat#HY-12299 |
| ZM-44739 | MedChemExpress | Cat#HY-10128 |
| **Commercial assays and reagents** | | |
| IncuCyte® Cytotox Green Reagent | Sartorius | Cat#4633 |
| **RNeasy blood midi kit** | Qiagen | Cat#52304 |
| U-PLEX Custom Biomarker Group 1 (human) Assays | Meso Scale Discovery | K15067M |
| U-PLEX Human TGF-β1 Assay | Meso Scale Discovery | K151XWK |
| V-PLEX Vascular Injury Panel 2 Human Kit | Meso Scale Discovery | K15198D |
| **Filters** | | |
| Cell strainer pore size 100 µM | Corning | Cat# CLS431752 |
