## Supplemental Table 3 for "Comparative characterization of OncoPro™ and Wnt-Based media reveals distinct phenotypic and pharmacologic states in patient-derived tumor organoids"

| Experiment | Organoid line | Passage number |
|---|---|---|
| RNA | PDAC136_WNT | 18 |
| RNA | PDAC136_OP | 22 |
| RNA | PDAC138_WNT | 22 |
| RNA | PDAC138_OP | 23 |
| RNA | PDAC154_WNT | 17 |
| RNA | PDAC154_OP | 19 |
| RNA | CRC013_WNT | 15 |
| RNA | CRC013_OP | 15 |
| RNA | CRC014_WNT | 7 |
| RNA | CRC014_OP | 6 |
| RNA | CRC018_WNT | 8 |
| RNA | CRC018_OP | 10 |
| RNA | CUPASC001_WNT | 12 |
| RNA | CUPASC001_OP | 13 |
| RNA | HNSCC007_WNT | 18 |
| RNA | HNSCC007_OP | 20 |
| RNA | GEJC025_WNT | 24 |
| RNA | GEJC025_OP | 24 |
| RNA | GEJC029_WNT | 19 |
| RNA | GEJC029_OP | 18 |
| RNA | CRC038_WNT | 16 |
| RNA | CRC038_OP | 17 |
| DNA | PDAC136_WNT | 18 |
| DNA | PDAC136_OP | 22 |
| DNA | PDAC138_WNT | 22 |
| DNA | PDAC138_OP | 23 |
| DNA | PDAC154_WNT | 17 |
| DNA | PDAC154_OP | 19 |
| DNA | CRC013_WNT | 15 |
| DNA | CRC013_OP | 15 |
| DNA | CRC014_WNT | 7 |
| DNA | CRC014_OP | 6 |
| DNA | CRC018_WNT | 8 |
| DNA | CRC018_OP | 10 |
| DNA | CUPASC001_WNT | 12 |
| DNA | CUPASC001_OP | 13 |
| DNA | HNSCC007_WNT | 18 |
| DNA | HNSCC007_OP | 20 |
| DNA | GEJC025_WNT | 24 |
| DNA | GEJC025_OP | 24 |
| DNA | GEJC029_WNT | 19 |
| DNA | GEJC029_OP | 18 |
| DNA | CRC038_WNT | 16 |

| | | |
|---|---|---|
| DNA | CRC038_OP | 17 |
| Drug screen biological rep 1 | PDAC138_WNT | 24 |
| Drug screen biological rep 2 | PDAC138_WNT | 24 |
| Drug screen biological rep 1 | PDAC138_OP | 24 |
| Drug screen biological rep 2 | PDAC138_OP | 24 |
| Drug screen biological rep 1 | PDAC154_WNT | 26 |
| Drug screen biological rep 2 | PDAC154_WNT | 26 |
| Drug screen biological rep 1 | PDAC154_OP | 41 |
| Drug screen biological rep 2 | PDAC154_OP | 41 |
| Drug screen biological rep 1 | CRC018_WNT | 19 |
| Drug screen biological rep 2 | CRC018_WNT | 19 |
| Drug screen biological rep 1 | CRC018_OP | 19 |
| Drug screen biological rep 2 | CRC018_OP | 19 |
| Drug screen biological rep 1 | CRC018_OP_WNT | 26 |
| Drug screen biological rep 2 | CRC018_OP_WNT | 26 |
| Drug screen biological rep 1 | PDAC138_WNT_OP_ | 21 |
| Drug screen biological rep 2 | PDAC138_WNT_OP_ | 21 |
| Cytokine assay | CRC018_WNT | 14 |
| Cytokine assay | CRC018_OP | 16 |
