## Supplemental Table 4 for "Comparative characterization of OncoPro™ and Wnt-Based media reveals distinct phenotypic and pharmacologic states in patient-derived tumor organoids"

| Cytokine (abbreviation) |
|---|
| BDNF |
| CRP |
| ctack |
| eotaxin |
| epo |
| fractalkine |
| g-csf |
| gm-csf |
| Ifn-β |
| ifn-γ |
| il-10 |
| il-12/il-23p40 |
| il-12p70 |
| il-13 |
| il-15 |
| il-16 |
| il-17a |
| il-17F |
| il-18 |
| il-1RA |
| il-1β |
| il-2 |
| il-2 Rα |
| il-21 |
| il-22 |
| il-23 |
| il-3 |
| il-33 |
| il-4 |
| il-5 |
| il-6 |
| il-7 |
| il-8 |
| il-9 |
| INF-a2a |
| ip-10 |

| |
|---|
| **mcp-1** |
| **mcp-2** |
| **MCP-3** |
| **m-csf** |
| **mip-1α** |
| **SAA** |
| **siCAM** |
| **sVCAM-1** |
| **tgf-b total** |
| **tgf-b_active** |
| **tnf-α** |
| **TSLP** |
| **mip-1β** |
| **mip-3α** |

| **Cytokine (full name)** |
|---|
| brain-derived neurotrophic factor |
| C-reactive proteine |
| Cutaneaous T-cell attracting chemokine |
| Eotaxin |
| Erythropoietin |
| Fraktalkine (CX3CL1) |
| Granulocyte- colony stimulating factor |
| granulocyte-macrophage- colony stimulating factor |
| Interferon beta |
| Interferon-gamma |
| Interleukin-10 |
| Interleukin-12/Interleukin-23p40 |
| Interleukin-12p70 |
| Interleukin-13 |
| Interleukin-15 |
| Interleukin-16 |
| Interleukin-17a |
| Interleukine 17F |
| Interleukin-18 |
| interleukine 1 receptor antagonist |
| Interleukin-1β |
| Interleukin-2 |
| Interleukine 2 Rα |
| Interleukine 21 |
| Interleukine 22 |
| Interleukine 23 |
| Interleukine 3 |
| Interleukine 33 |
| Interleukin-4 |
| Interleukin-5 |
| Interleukin-6 |
| Interleukin-7 |
| Interleukin-8 |
| Interleukine 9 |
| Interferon a2a |
| Interferon-gamma induced protein 10 |

| |
|---|
| Monocyte-chemoattractant protein 1 |
| Monocyte-chemoattractant protein 2 |
| Monocyte-chemoattractant protein 3 |
| Macrophage-colony stimulating factor |
| Macrophage inflammatory protein-1 alpha |
| Serum amyloid A |
| soluble intercellular adhesion molecule 1 |
| soluble vascular cell adhesion molecule 1 |
| Transforming growht factor beta (total) |
| Transforming growht factor beta (only active form) |
| Tumor necrosis factor alpha |
| Thymic stromal lymphopoietin |
| Macrophage inflammatory protein-1 beta |
| Macrophage inflammatory protein-3 alpha |
