## Supplemental Table 5 for "Comparative characterization of OncoPro™ and Wnt-Based media reveals distinct phenotypic and pharmacologic states in patient-derived tumor organoids"

| Organoid name | Sex | Tumor location | Tumor type |
|---|---|---|---|
| **CCA_012** | F | CCA | Adenocarcinoma |
| **CCA_ASC_003** | M | CCA | Adenocarcinoma |
| **CRC_009** | M | CRC | Adenocarcinoma |
| **CRC_013** | F | CRC | Adenocarcinoma |
| **CRC_014** | F | CRC | Adenocarcinoma |
| **CRC_018** | M | CRC | Adenocarcinoma |
| **CUP_ASC_001** | F | CUP | Adenocarcinoma |
| **GEJC_025** | M | GEJC | Adenocarcinoma |
| **GEJC_029** | M | GEJC | Adenocarcinoma |
| **CRC_038** | F | CRC | Adenocarcinoma |
| **HNSCC_001** | M | HNSCC | Spinocellular carcinoma |
| **HNSCC_002** | M | HNSCC | Spinocellular carcinoma |
| **HNSCC_007** | M | HNSCC | Spinocellular carcinoma |
| **PDAC_044** | F | PDAC | Adenocarcinoma |
| **PDAC_136** | M | PDAC | Adenocarcinoma |
| **PDAC_138** | M | PDAC | Adenocarcinoma |
| **PDAC_154** | F | PDAC | Adenocarcinoma |
| **BR_007** | F | Breast cancer | Adenocarcinoma |
| **BR_008** | F | Breast cancer | Adenocarcinoma |
| **CCA_ASC_002** | F | CCA | Adenocarcinoma |
| **CRC_019** | F | CRC | Adenocarcinoma |
| **CRC_020** | M | CRC | Adenocarcinoma |
| **CRC_023** | M | CRC | Adenocarcinoma |
| **GI_ASC_001** | M | Gastric | Adenocarcinoma |
| **PDAC_060** | M | PDAC | Adenocarcinoma |
| **PDAC_087** | M | PDAC | Adenocarcinoma |
| **PDAC_164** | F | CCA | Adenocarcinoma |
| **PDAC_165** | M | PDAC | Adenocarcinoma |
| **PDAC_167** | M | PDAC | Adenocarcinoma |
| **PDAC_168** | F | PDAC | Adenocarcinoma |
| **PDAC_169** | M | Renal cell carcinoma | Adenocarcinoma |
| **PDAC_170** | M | PDAC | Adenocarcinoma |
| **PDAC_173** | M | PDAC | Adenocarcinoma |
| **PDAC_178** | F | PDAC | Adenocarcinoma |
| **Lu_02_021** | M | NSCLC | Spinocellular carcinoma |
| **Lu_02_023** | F | NSCLC | Adenocarcinoma |

| Pretreated? | Preatreatment Regimen | TNM classification | Differentiation g | Microsatellite s |
|---|---|---|---|---|
| Yes | Gemcitabine/cisplatinum; FOLFOX | ypT3N0M0 | Poorly | MSS |
| Yes | Gemcitabine/cisplatine/durvaluma | cTxNxM1 | Unknown | MSI |
| No | NA | pT3N0M0 | Moderately | MSS |
| Yes | Neoadjuvant nivolumab + RT | ypT3N2M0 | Moderately | MSS |
| No | NA | pT3N0M0 | Moderately | MSI |
| No | NA | pTxNxM1 | Moderately | MSS |
| Yes | Carboplatin-paclitaxel-bevacizuma | pTxNxM1 | Unknown | MSS |
| Yes | CROSS | ypT3N1M0 | Poorly | Unknown |
| No | NA | pT3N0M0 | Poorly | MSS |
| No | NA | pT3N1M0 | Moderately | MSS |
| No | NA | pT2N1M0 | Unknown | Unknown |
| No | NA | pT1aN0M0 | Unknown | Unknown |
| No | NA | cT2N1M0 | Unknown | Unknown |
| Yes | Gemcitabine/nab-paclitaxel | ypT3N2M0 | Poorly | Unknown |
| No | NA | pT3N1M0 | Moderately | Unknown |
| No | NA | pT3N2M0 | Poorly | Unknown |
| No | NA | pT2N2M0 | Poorly | MSS |
| No | NA | pT4bN1aM0 | Poorly | Unknown |
| No | NA | pT2N0M0 | Poorly | Unknown |
| Yes | Gemcitabine/cisplatinum/durvalum | cTxNxM1 | Unknown | Unknown |
| No | NA | pT3N1M0 | Moderately | MSS |
| No | NA | pT3N2aMx | Moderately | MSS |
| No | NA | pT2N0M0 | Moderately | Unknown |
| Yes | FLOT; FOLFIRI/ramucirumab | cT4NxM1 | Unknown | MSS |
| No | NA | pT2N1M0 | Moderately-poor | MSS |
| Yes | FOLFIRINOX | ypT3N1M0 | Moderately | Unknown |
| No | NA | pT3N2M0 | Poorly | Unknown |
| No | NA | pT3N2M0 | Moderately | Unknown |
| No | NA | pT2N2M0 | Poorly | Unknown |
| No | NA | pT4N0M0 | Moderately | Unknown |
| No | NA | pTxNxM1 | ISUP grade 2 | Unknown |
| Yes | FOLFIRINOX | ypT2N2M1 | Moderately | Unknown |
| No | NA | pT2N1Mx□ | Poorly | Unknown |
| No | NA | pT2N1M0 | Poorly | Unknown |
| Yes | Gemcitabine/cisplatine/pembrolizu | ypT2N1M0 | Unknown | Unknown |
| No | NA | pT3N0M0 | Grade 2 | Unknown |

| Mutations | Comment |
|---|---|
| KRASG13D | |
| BRAF non-V600E | Ascites fluid; isolated loss of MSH6 |
| Unknown | concurrent lymphoma in colon; rectal |
| Unknown | rectum |
| Unknown | Right colon |
| Unknown | From lung metastasis; rectum |
| KRASG12R | Ascites fluid |
| Unknown | |
| Unknown | FISH HER2+ |
| Unknown | Left colon |
| Unknown | Oral cavity; p16+ |
| Unknown | Sinus piriformis; p16- |
| Unknown | Hypopharynx |
| Unknown | |
| Unknown | |
| KRASG12D | |
| Unknown | |
| Unknown | |
| Unknown | ER/PR positive; concurrent NSCLC |
| IDH1 | Triple negative breast cancer; Ascites fluid |
| Unknown | Right colon |
| Unknown | Left colon |
| Unknown | rectum |
| Unknown | Ascites fluid; diffuse type |
| KRASG12A | |
| Unknown | |
| Unknown | |
| Unknown | |
| Unknown | |
| Unknown | concurrent transitional cell cancer |
| Unknown | |
| Unknown | |
| Unknown | |
| Unknown | concurrent NSCLC |
| Unknown | |
| KRASG12V | non-mucinous |
